# The *E. coli* DnaX clamp loader sharply bends DNA to load β-clamp at nicks and small gaps

**DOI:** 10.64898/2026.01.17.700081

**Authors:** Fengwei Zheng, Nina Y. Yao, Roxana E. Georgescu, Meinan Lyu, Michael E. O’Donnell, Huilin Li

**Author notes:** These authors contributed equally to this work (Fengwei Zheng, Nina Y. Yao). Correspondence should be addressed to M.E.O or H.L.

## Abstract

DNA sliding clamps are essential for processive DNA synthesis in all domains of life and are loaded by ATP-dependent clamp loaders that recognize recessed 3′ ends. How clamp loaders function at nicks and small ssDNA gaps—common intermediates during DNA repair—remains incompletely understood. Here, we show that the bacterial *Escherichia coli* DnaX clamp loader employs a fundamentally different mechanism from its eukaryotic counterpart. Whereas eukaryotic RFC unwinds DNA at the recessed 3′ end and stabilizes the 5′-dsDNA at a dedicated shoulder site, the bacterial DnaX-complex neither unwinds DNA nor stably binds the 5′-dsDNA in vitro. Instead, cryo-EM structures reveal that the β-clamp itself contains a conserved external DNA-binding site that enables sharp bending of gapped DNA by ∼150°, promoting insertion of the 3′-dsDNA into the clamp. This DNA-bending mechanism allows efficient β-clamp loading at nicks and small gaps and reveals a distinct bacterial strategy for clamp loading. Because small DNA gaps are frequently associated with DNA damage, clamps loaded at these sites are likely important for DNA repair.

**In brief:** Zheng et al. show that the bacterial clamp loader DnaX-complex uses a DNA-bending mechanism—rather than DNA unwinding—to load the β-clamp at nicks and small gaps, revealing a clamp-loading strategy distinct from eukaryotic RFC and relevant to DNA damage repair.

**Highlights:**

- The bacterial DnaX clamp loader lacks a stable shoulder DNA-binding site
- Unlike eukaryotic RFC, DnaX does not unwind DNA at nicks and small gaps
- The E. coli β-clamp contains a conserved external DNA-binding site absent in PCNA
- Sharp DNA bending enables β-clamp loading at nicks and small ssDNA gaps

## Introduction

In all cell types—bacteria, archaea and eukaryotes—the DNA replication machinery utilizes ring-shaped clamps that encircle duplex DNA and tether replicative DNA polymerases (Pols) to DNA, enabling rapid and highly processive synthesis^1–4^. This function is readily apparent in the *Escherichia coli* (*E. coli*) Pol III replication system, in which circular clamps were first discovered^1^. In the absence of a clamp, *E. coli* Pol III core synthesizes DNA at only ∼1 nucleotide (nt) per second; upon engagement with its β-clamp, Pol III synthesizes DNA at >500 nt per second with extremely high processivity (>5 kb)^1^. Because circular sliding clamps cannot self-assemble onto DNA, they require an ATP-driven clamp loader that recognizes a DNA 3′-recessed end and loads the clamp onto it^3,4^.

In *E. coli*, the β-clamp is a homodimer, with each protomer containing three domains of identical fold, giving the clamp pseudo–six-fold symmetry. The eukaryotic equivalent, PCNA, is a homotrimer in which each subunit contains two domains of the same fold, also forming a pseudo–six-fold symmetric ring^2,5,6^ (**Figure 1A**). The *E. coli* clamp loader motor consists of one δ subunit, one δ′ subunit, and three *dnaX*-encoded subunits^7,8^. The *dnaX* gene produces two isoforms: a full-length τ subunit and a shorter γ subunit generated by translational frameshifting^8^ (**Figure 1B**). In this study, we refer to this pentameric motor assembly as the DnaX-complex. The bacterial clamp loader also contains two small accessory subunits, χ and ψ, which are not required for clamp loading per se^8^. The full seven-subunit assembly is referred to here as the DnaX-holo-complex. The eukaryotic counterpart of the DnaX-complex is the heteropentameric Replication Factor C (RFC) clamp loader^4,9^.

**Figure 1.**
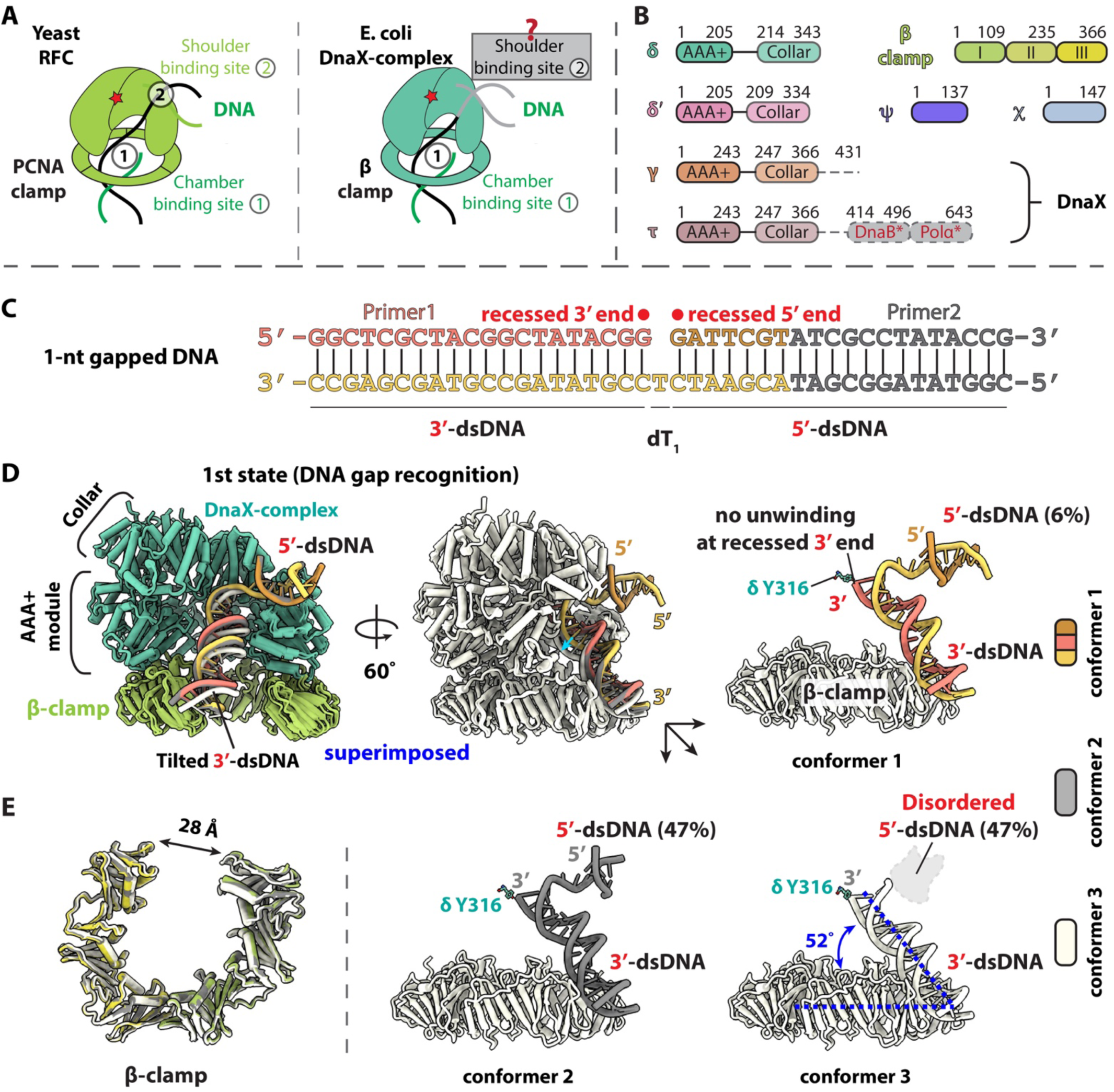
The *E. coli* DnaX-complex loading β-clamp onto nicked or 1-nt gapped DNA. (**A**) Schematic illustrating that eukaryotic RFC uses a shoulder DNA-binding site to facilitate PCNA loading onto small-gaped DNA. In contrast, how the bacterial DnaX-complex loads the β-clamp onto nicked or small gapped DNA, and whether it possesses a functional shoulder DNA-binding site, has remained unclear. (**B**) Domain organizations of the DnaX-holo-complex and the β-clamp. The *E. coli dnaX* gene encodes the full-length τ subunit and a C-terminally truncated γ subunit lacking the C-terminal domain (CTD). The mobile τ CTD (grey) contains regions that interact with DnaB helicase and Pol III α (asterisks). (**C**) Sequence of the 1-nt gapped DNA substrate used for cryo-EM study. The unresolved region is shown in gray. The duplex DNA segment containing the recessed 3′ end is referred to as the 3′-dsDNA, and the duplex segment containing the recessed 5′ end is referred to as the 5′-dsDNA. (**D**) Three conformers of the first major loading state—the DNA recognition state—of the DnaX-complex–β-clamp bound to 1-nt gapped DNA. The top two panels show two views of the three superimposed conformers. In all conformers, the recessed DNA 3′ end has entered the clamp loader chamber, whereas the 3′-dsDNA remains outside the β-clamp. The 5′-dsDNA is partially ordered in conformer 1 (7 bp) and conformer 2 (5 bp) but is completely disordered in conformer 3 (bottom panels). Notably, no base pair separation is observed at the recessed 3′ end, indicating that the δ-subunit Y316 does not unwind DNA (lower left panel). (**E**) Superposition of the β-clamps from the three conformers showing the identical gate opening of ∼28 Å. Unless otherwise indicated, distance measurements were performed using surface representation in ChimeraX^75^.

Crystal structures of both the DnaX-complex and RFC revealed that these clamp loaders adopt similar overall architectures^7,10^. Each forms a spiral-shaped pentamer with a central chamber that accommodates dsDNA and features a prominent gap between δ and δ′ subunits through which DNA enters the chamber. The sliding clamp is positioned directly beneath this chamber, explaining how clamp loaders guide DNA into the open clamp^3^. In both systems, ATP-binding sites are located at subunit interfaces, a hallmark of AAA+ ATPases, allowing coordinated inter-subunit communication during the clamp loading cycle^3,11^.

Clamp loading at large ssDNA gaps, such as those between Okazaki fragments, is well understood for both eukaryotic RFC^12–15^ and the bacterial DnaX-complex^16,17^. In both cases, the clamp remains closed in solution until binding by the clamp loader induces gate opening^18^. A notable difference is that DnaX-complex opens the β-clamp in a counterclockwise direction, whereas RFC opens PCNA clockwise^16^, consistent with earlier biochemical studies demonstrating opposite directionality in bacterial and eukaryotic clamp opening^19,20^. Following clamp opening, the DNA 3′-recessed end is recognized by the clamp loader chamber, dsDNA enters the clamp, and the clamp subsequently closes around the DNA. These steps require ATP binding but not ATP hydrolysis^16,17,21^. ATP hydrolysis then triggers dissociation of the clamp loader, completing the loading reaction^22–24^.

Biochemical studies have shown that both the bacterial and eukaryotic clamp loaders can load clamps at a nick or small ssDNA gaps (<6 nt), a process relevant to DNA repair. Clamp loading at such sites is more challenging and remains an active area of investigation. Recent structural studies demonstrated that eukaryotic RFC unwinds DNA from the 3′-recessed end to widen small gaps and revealed an external “shoulder” DNA-binding site that stabilizes the dsDNA segment at the 5′-recessed end (hereafter 5′-dsDNA)^12–15^. This shoulder site is formed by the unique Rfc1 BRCT domain together with the Rfc1 AAA+ domain and allows RFC to simultaneously bind both the 3′- and 5′-dsDNA segments^25–27^ (**Figure 1A**). The ability of RFC to function at small gaps has been shown to correlate with DNA repair^12–15^.

How the bacterial DnaX-complex loads clamps onto nicked or small-gapped DNA has not previously been visualized. Consequently, it has remained unclear whether the bacterial clamp loader possesses a functional shoulder DNA-binding site or whether it unwinds DNA in a manner analogous to eukaryotic RFC (**Figure 1A**). To address these questions, we used cryo-EM to determine structures of the DnaX-complex bound to the β-clamp and DNA substrates containing 1- or 10-nt gaps. We captured a total of 13 clamp loading intermediates (**Supplemental video 1**) and found that, in contrast to RFC, the bacterial DnaX-complex neither unwinds DNA nor stably binds the 5′-dsDNA at the shoulder site under our in vitro conditions. Instead, we uncover a distinct clamp loading mechanism in which the β-clamp utilizes an external DNA-binding site to enable sharp DNA bending, allowing clamp loading at nicks and small gaps.

## RESULTS

### 1. Cryo-EM of bacterial DnaX-complex with β-clamp and 1-nt gapped DNA

Previous studies of the eukaryotic RFC demonstrated that it can unwind up to 6 bp at a DNA nick to generate a sufficiently long ssDNA gap for clamp loading^14,15^. Although earlier biochemical studies showed that the *E. coli* DnaX-complex can load clamps onto nicked DNA^1^, whether the bacterial loader unwinds DNA in a manner analogous to eukaryotic RFC has remained unclear (**Figure 1A**).

The *E. coli dnaX* gene encodes two isoforms, the full-length τ subunit and a C-terminally truncated γ subunit. As a result, the seven-subunit bacterial loader complex can exist in two forms, one containing three τ subunits (τ_3_δδ′) and another containing three γ subunits (γ_3_δδ′) (**Figure 1B**). In this study, we used the full-length τ-containing complex (τ_3_, hereafter referred to as “τ-complex”) for cryo-EM analysis to capture structural features of the τ C-terminal region that was absent in previous crystal structures^7,28^. For subsequent biochemical assays, we used the γ-containing complex (γ_3_, hereafter referred to as the“γ-complex”), which has equivalent clamp loading activity and has been extensively characterized in prior biochemical studies^29,30^.

To investigate how the DnaX-complex loads the β-clamp onto small gaps (<6 nt) and whether it unwinds DNA in this context, we assembled in vitro clamp loading intermediates by mixing purified τ-complex, β-clamp, and a 1-nt gapped DNA substrate (**Figure 1C**) at a molar ratio of 1:1.3:1.5 in the presence of 5 mM Mg^2+^ and 0.5 mM ATPγS, a slowly hydrolyzable ATP analog^31^. Cryo-EM grids were prepared from this mixture, and data were collected and processed as described below. Two major loading states were resolved, corresponding to early and late stages of the clamp loading reactions (**Figure S1**, **Table S1**).

### 2. DnaX-complex does not possess a stable shoulder binding site for 5′-dsDNA

In the first major loading state, the β-clamp is open, but the 3′-dsDNA has yet entered the clamp. The DNA 3′-recessed end resides inside the loader chamber, oriented towards the δ-subunit residue Y316. However, the 3′-dsDNA tilts outwards by ∼52° away from the β-clamp and does not enter the clamp interior, but instead rests on the outer surface of the open clamp. We therefore refer to this conformation as the “DNA gap recognition” state (**Figure 1D**). This disposition of the 3′-dsDNA is unprecedented. In all structurally visualized clamp loading intermediates—whether bacterial or eukaryotic—the 3′-dsDNA is either unresolved or has fully entered the clamp chamber.

Density corresponding to the 5′-dsDNA was weak and variable near the shoulder region of the DnaX-complex (**Figure 1D**). Further 3D classification resolved three conformers at resolutions ranging from 3.14 to 2.71 Å, differing primarily in the extent of 5′-dsDNA ordering (**Figure S1**). In conformer 1, approximately 9 bp of the 5′-dsDNA are ordered; in conformer 2, only 5 bp are ordered; and in conformer 3, the 5′-dsDNA is completely disordered. Notably, conformer 1 accounts for only ∼6% of particles in this state, indicating that the DnaX-complex does not stably bind the 5′-dsDNA at the shoulder site. Moreover, from conformer 1 to conformer 3, the 3′-dsDNA moves incrementally toward clamp chamber, as indicated by the cyan arrow in **Figure 1D**.

In this DNA gap recognition state, the β-clamp gate is opened by ∼28 Å (**Figure 1E**), which is wider than the 20 Å opening previously observed for the DnaX-complex bound to a tailed DNA substrate^16^. This wider opening may be stabilized by interactions with the blunt end of the 3′-dsDNA.

### 3. DnaX-complex sharply bends DNA for loading β-clamp onto 1-nt gapped DNA

In the second major loading state, the 3′-dsDNA has fully entered the chambers of both the DnaX-complex and the β-clamp, representing a post–DNA–entry state. Further classification of this state yielded two conformers at resolutions of 2.72 Å and 3.12 Å, respectively, which differ in β-clamp gate width and DNA positioning (**Figures 2A, S1**).

**Figure 2.**
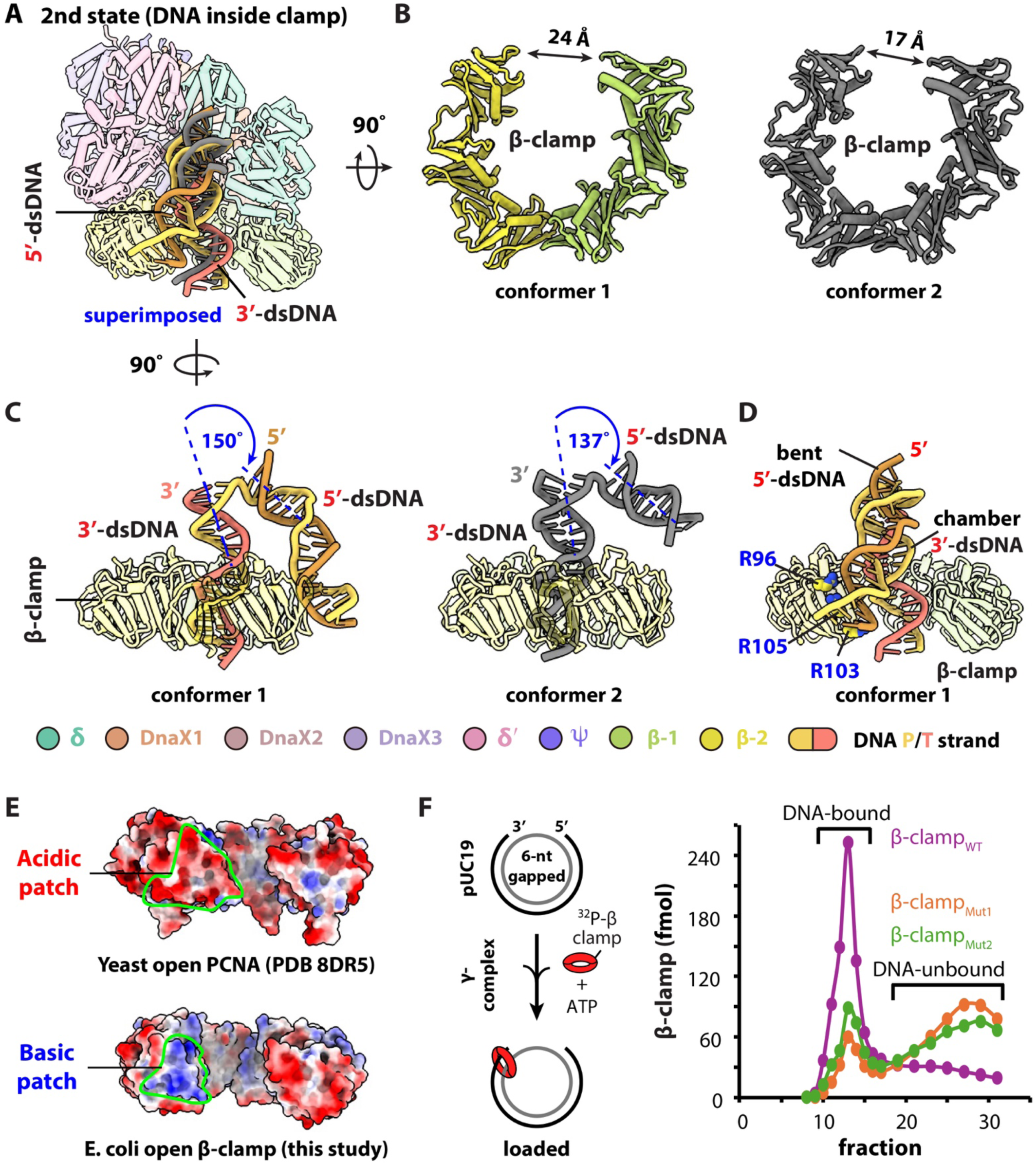
The second loading state of the DnaX-complex during β-clamp loading onto 1-nt gapped DNA and the role of a β-clamp basic patch. (**A**) Superimposition of conformers 1 (color) and 2 (gray) of the second major loading state, in which the 3′-dsDNA has entered the β-clamp chamber. (**B**) Side-by-side comparison of the β-clamp in conformers 1 and 2, highlighting differences in clamp gate width. (**C**) Side view of the superimposed structures in (A), with the DnaX-complex omitted for clarity. The two conformers are shown separately. In conformer 1, the 5′-dsDNA bends sharply and binds the outer surface of the β-clamp at the site previously occupied by the 3′-dsDNA. In conformer 2, 5′-dsDNA dissociates from the β-clamp and becomes partially flexible (11 bp ordered), concomitant with further clamp closure. (**D**) Three positively charged residues (R96, R103, and R105) form a distinct basic patch on the outer surface of the β-clamp near the left gate. (**E**) Comparison of this region with yeast PCNA reveals that the corresponding surface is acidic. (**F**) Substitution of the three arginine residues with alanine (β-clamp_Mut1_) or glutamate (β-clamp_Mut2_) reduced β-clamp loading efficiency onto a 6-nt gapped pUC19 plasmid DNA to approximately one-third of that observed with wild-type β-clamp (β-clamp_WT_). In panels (**A)**, (**C**), and (**D**), protein components are shown as 80% transparent cartoons for clarity.

In state 2 conformer 1, the β-clamp gate narrows from the 28 Å opening observed in the DNA gap recognition state to 24 Å. In this conformer, the 5′-dsDNA bends sharply downward by ∼150° and binds to the outer surface of the β-clamp at the site previously occupied by the 3′-dsDNA in the DNA gap recognition state (**Figure 2A**−**C**). In state 2 conformer 2, the 3′-dsDNA enters the β-clamp further, the 5′-dsDNA dissociates from the clamp and became less sharply bent (∼137°), and the β-clamp gate narrows further to ∼17 Å (**Figure 2B**−**C**).

Together, these two conformers suggest a clamp-loading mechanism in which sharp bending of the 5′-dsDNA promotes insertion of the 3′-dsDNA into the β-clamp chamber, followed by progressive clamp closure.

### 4. A unique basic patch on β-clamp is utilized in clamp loading onto small gapped DNA

The repeated binding of 3′-dsDNA in the first loading state and 5′-dsDNA in the second loading state binding to the outer surface of the open β-clamp led us to identify a basic patch on the β-clamp surface. This patch is formed by three positively charged residues—R96, R103, and R105—which lie within ∼3.5 Å of the DNA (**Figure 2D**). Structure-based sequence alignment of bacterial β-clamps, spanning sequence identities 29% to 96%, revealed strong conservation of those three arginine residues (**Figure S2A–B**). In contrast, the corresponding region in yeast and human PCNA is acidic (**Figure 2E**).

To examine the functional importance of this basic patch, we incorporated an N-terminal peptide ^32^P-labeling motif in wild-type β-clamp (β-clamp_WT_) and generated two mutants by substituting the arginine residues with either alanine (β-clamp_Mut1_) or aspartic acid (β-clamp_Mut2_). We then quantified clamp loading efficiency onto pUC19 plasmid DNA containing a 6-nt gap using a gel-filtration–based assay^19,32–34^ (see **STAR METHODS**). Consistent with previous studies indicating that gaps of ≥6 nt are optimal for clamp loading^13–15,31^, both β-clamp_Mut1_ and β-clamp_Mut2_ exhibited markedly reduced loading efficiencies, achieving ∼35% and ∼25%, respectively, of the loading observed with β-clamp_WT_ (**Figure 2F**). Mass photometry confirmed that both mutants remained dimeric in solution (**Figure S2C**),

We further examined the ability of these β-clamp basic patch mutants to assemble with the DnaX-complex and 1-nt gapped DNA by cryo-EM. 2D classification and 3D reconstruction revealed that, in contrast to wild-type β-clamp, the mutant clamps failed to form complexes in which DNA binds directly to β-clamp surface (**Figure S3A-B**). Together, these results demonstrate a functional role for the conserved β-clamp basic patch in DNA binding and clamp loading at small gaps.

In addition, we note that a recent cryo-EM study by Landeck et al. reported that, in the absence of DNA, β-clamp domain I—which harbors the basic patch—is disordered^17^. In a comparable state observed here (RMSD_Cα_ 0.9 Å), binding of DNA stabilizes this domain, indicating that DNA engagement at the basic patch contributes to the ordering of β-clamp domain I (**Figure S3C**).

### 5. Y316 recognizes but does not separate the DNA 3′ end

The overall architectures of the three conformers in the first loading state and the two conformers in the second loading state are highly similar. Superimposition of the first four conformers onto state 2 conformer 2 yields RMSD_Cα_ values ranging from 0.8 to 1.0 Å, indicating that the primary structural differences among these intermediates reside in the DNA rather than in the protein components (**Figures 1C and 2A**).

Closer examination of residue Y316 of the δ-subunit, previously termed a “separation pin” ^28^, revealed its role in recognizing—but not unwinding—the DNA 3′ end. In the first loading state, before the 3′-dsDNA enters the loader chamber, Y316 is only partially ordered. From conformer 1 to conformer 3, the distance between the aromatic ring of Y316 and the DNA 3′-junction (i.e., last base of primer 1, G20) decreases from 12.0 Å to 11.4 Å and then to 9.9 Å (**Figure S4**, left). Concurrently, the local cryo-EM density for Y316 becomes progressively stronger, consistent with increasing stabilization of this residue.

In the second loading state, after the 3’-dsDNA has entered the chamber, Y316 becomes fully ordered and adopts a conformation that mimics a DNA base, stacking against the terminal base of the primer 3’ end. In this state, the distance between Y316 and the DNA 3’ end is further reduced to 3.5 Å and 3.1 Å in conformers 1 and 2, respectively (**Figure S4**, right). Across all five conformers, we did not observe separation or unwinding of the DNA strands at the 3′ end.

This absence of DNA unwinding contrasts with observations in eukaryotic RFC clamp loaders, in which the analogous residue can unwind up to 6 bp at the DNA 3′ end^13–15,35^. Taken together, these findings indicate that, rather than functioning as a strand-separation element, Y316 promotes recognition and positioning of DNA 3’ end during gap sensing and clamp loading (**Figure S4**).

### 6. The efficiency with which the DnaX-complex loads the β-clamp onto DNA with nicks or small gaps

To test the hypothesis that the *E. coli* DnaX-complex employs a distinct strategy to load the β-clamp onto DNA containing nick or small gaps (<6 nt), we compared the clamp loading activities of the wild-type clamp loader (γ-complex_WT_) and a separation-pin mutant bearing the Y316A substitution (γ-complex_Mut_). Clamp loading was assayed using pUC19 plasmid DNA substrate containing either a nick or gaps of defined sizes (**Figure 3A**, see **Methods**).

**Figure 3.**
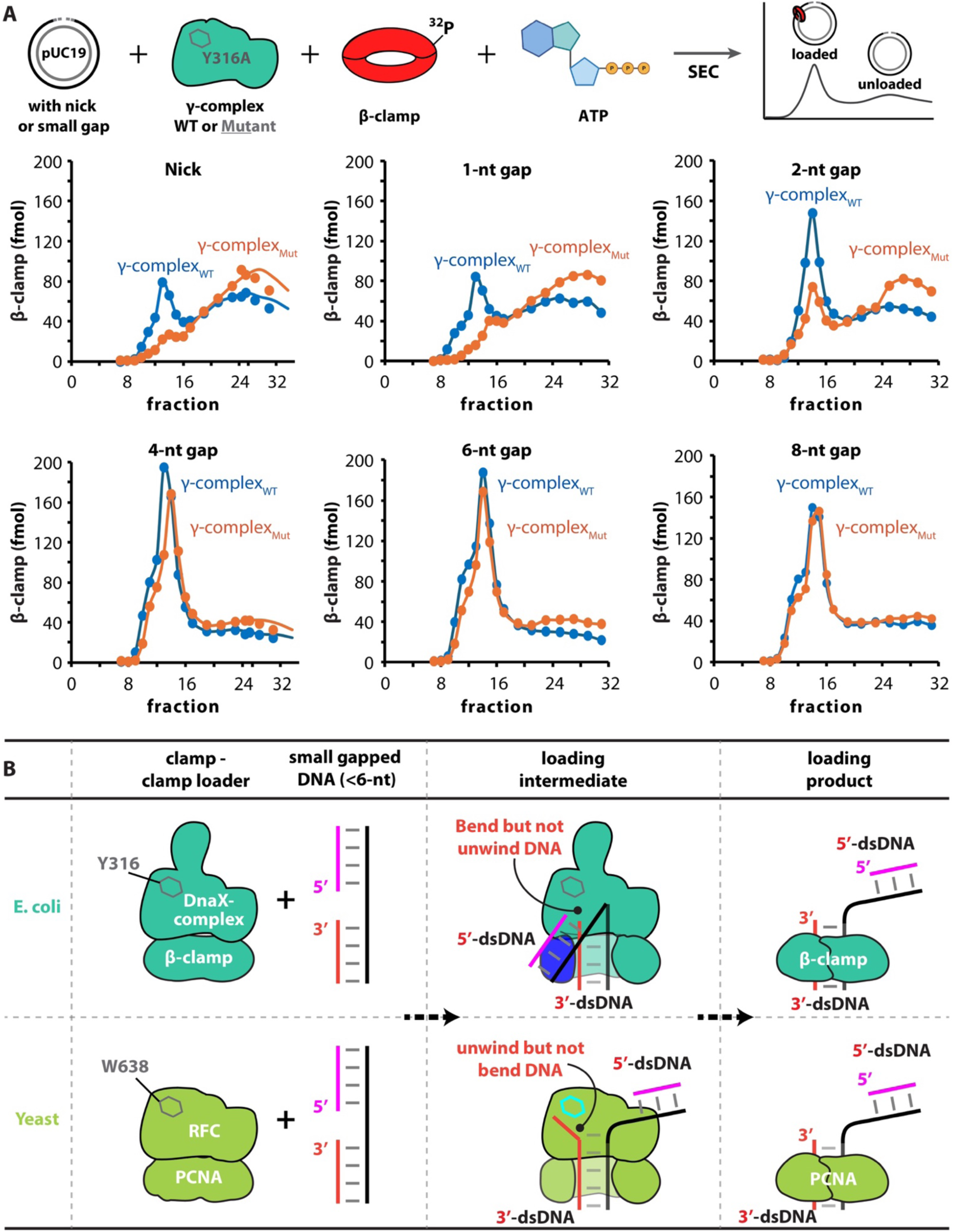
The *E. coli* DnaX-complex employs a DNA-bending strategy to load the β-clamp onto DNA containing small gaps (<6 nt). (**A**) Comparison of β-clamp loading efficiencies mediated by the wild-type clamp loader (γ-complex_WT_) and a separation-pin mutant harboring the δ-subunit Y316A substitution (γ-complex_Mut_) using plasmid DNA substrates containing a nick or gaps of defined sizes. From the γ-complex_Mut_, loading efficiency increases progressively with gap size, approaching that of the wild-type complex, reaching ∼85% efficiency on 4-nt gapped DNA and ∼90% on 6-nt gapped DNA (see **Methods** for details). (**B**) Schematic comparison of clamp loading mechanisms in bacteria (e.g., *E. coli*) and eukaryotes (e.g., *S. cerevisiae*) at nicked or small gapped DNA. In yeast, the Rfc1 separation pin residue W638 unwinds up to ∼6 bp at the recessed 3′ end to generate a sufficiently long gap for PCNA loading^14,15^. In contrast, the corresponding residue in *E. coli*, δ Y316, recognizes the recessed DNA 3′ end without unwinding DNA. Instead, the bacterial DnaX-complex, but not RFC, induces sharp DNA bending (∼150°) to enable clamp loading.

For the wild-type clamp loader, the loading efficiency of β-clamp_WT_ onto nicked and 1-nt gapped DNA was approximately half of that observed for DNA substrates containing gaps of 2 nt or larger. In contrast, the γ-complex_Mut_ exhibited a pronounced deficiency in loading the β-clamp onto nicked DNA as well as DNA substrates containing 1-nt and 2-nt gaps (**Figure 3A**, upper). However, loading efficiency by the γ-complex_Mut_ increased progressively with gap size, reaching ∼85% and ∼90% of wild-type level on DNA substrates containing 4-nt and 6-nt gaps, respectively, and becoming indistinguishable from wild-type on an 8-nt gapped DNA substrate (**Figure 3A**, lower).

To further correlate these biochemical results with structural observations, we performed cryo-EM analysis of β-clamp loading onto 1-nt gapped DNA using γ-complex_Mut_. Consistent with the loading assay results, we were able to capture only the clamp–clamp loader complex without bound DNA (**Figure S3**). Together, these biochemical and structural data are mutually consistent and strongly support the clamp loading model proposed in **Figure 3B**.

For loading of PCNA onto small-gapped DNA, the eukaryotic RFC clamp loader unwinds DNA from the 3′ end to widen the gap and stabilizes both the 3′- and 5′-dsDNA segments. In contrast, the bacterial DnaX-complex neither unwinds DNA nor stably binds the 5′-dsDNA at a shoulder site under our in vitro conditions. Instead, it loads the β-clamp onto DNA substrates containing gaps smaller than 6 nt by sharply bending the 5′-dsDNA by up to ∼150°, whereas RFC induces substantially less 5′-dsDNA bending (∼75°)^12–14^. To our knowledge, such an extent of 5′-dsDNA bending during clamp loading has not been previously observed (**Figure 3B; Supplemental Video 2**).

### 7. Cryo-EM of β-clamp loading intermediates with a 10-nt gapped DNA

Using the 1-nt gapped DNA substrate, we established that the *E. coli* DnaX-complex does not unwind the DNA 3′ end, in contrast to its eukaryotic counterpart, and that binding of the 5′-dsDNA at the shoulder site is transient. One possible explanation for these observations is that the extremely short gap physically constrains the DNA such that, when the 3′-dsDNA enters the loader chamber, the 5′-dsDNA is forced away from the shoulder. Alternatively, the bacterial clamp loader may simply lack a functional shoulder DNA-binding site. In addition, the previously unrecognized DNA-binding interaction with the β-clamp basic patch could, in principle, be specific to very short gaps. To distinguish between these possibilities, we performed cryo-EM analysis of β-clamp loading intermediates using a DNA substrate containing a 10-nt gap, which is sufficient long to allow independent positioning of the 3′- and 5′-dsDNA segments (**Figure 4A-B)**.

**Figure 4.**
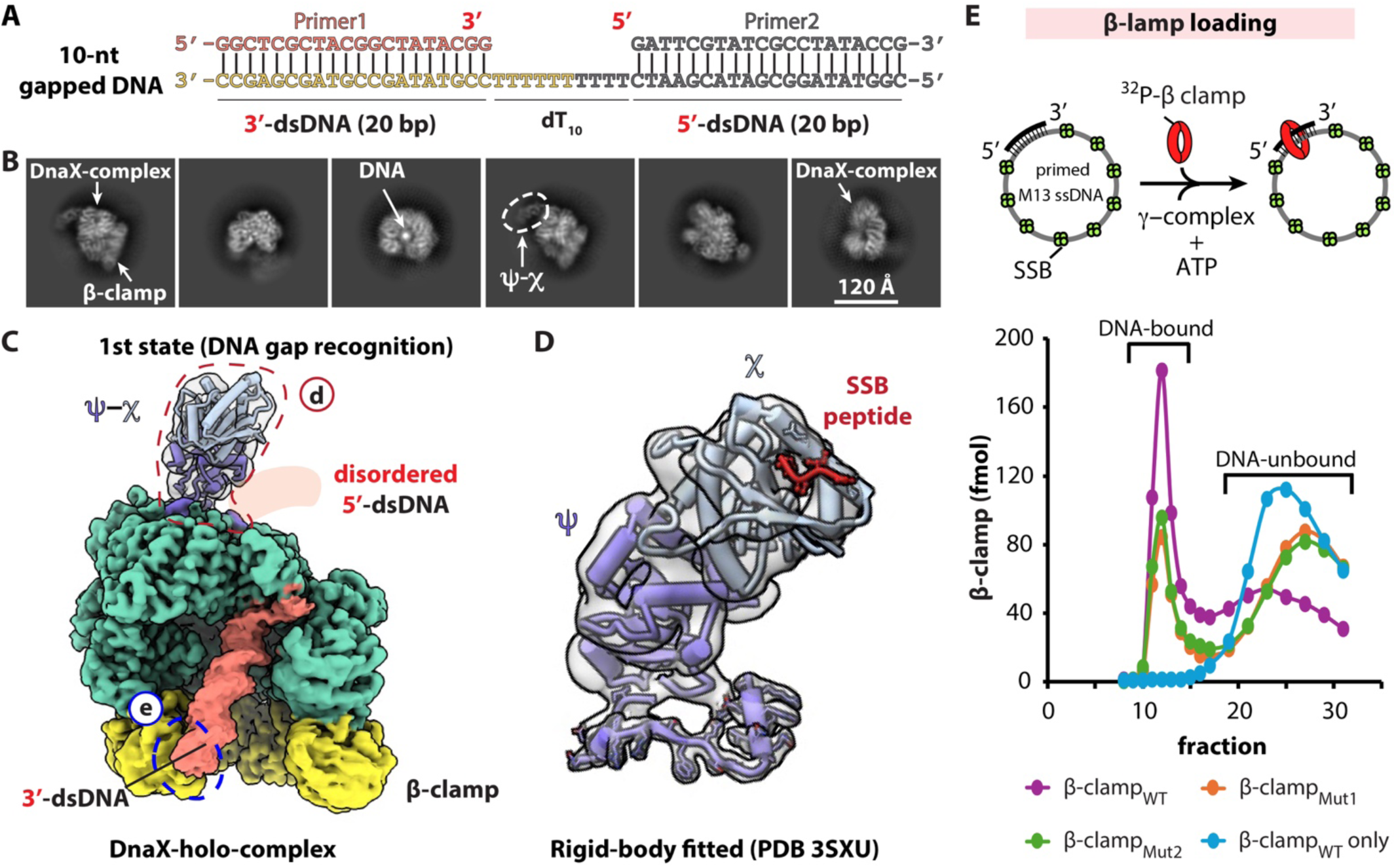
β-clamp loading by the DnaX-holo-complex onto a 10-nt gapped DNA substrate. (**A**) Architecture of the 10-nt gapped DNA used for cryo-EM analysis. (**B**) Representative 2D class averages of clamp loading intermediates. The dashed oval indicates weak density corresponding to the χψ heterodimer. (**C-D**) Front views of a DnaX-holo-complex in the 1st DNA gap recognition state during loading of the β-clamp onto a 10-nt gapped DNA. In this state, the 5′-dsDNA is disordered, whereas the 3′-dsDNA is ordered and bind to the outer surface of the β-clamp. The region outlined by the red dashed line (C) is enlarged in (**D**), showing a rigid-body docked χψ crystal structure bound to a SSB peptide^36^. The 3′-dsDNA is observed to bind the β-clamp basic patch (blue dashed outline), similar to that observed with 1-nt gapped DNA. (**E**) Loading efficiency of the β-clamp basic patch mutants on a singly primed DNA with a larger gap size. β-clamp_Mut1_ and β-clamp_Mut2_ harbor R96, R103, and R105 substitutions to alanine or glutamate, respectively, and were ^32^P-labelled for quantification alongside wild-type β-clamp (β-clamp_WT_).

Cryo-EM image processing yielded eight 3D cryo-EM maps at resolutions ranging from 2.95 to 2.53 Å, corresponding to eight distinct clamp-loading intermediates (**Figure S5, Tables S1** and **S2**). Although the full-length τ-containing DnaX-holo-complex was used for these structural studies, none of the resulting maps resolved density for the C-terminal domains of the three τ subunits, consistent with previous reports^16,17^. Accordingly, the atomic models built for these intermediates include only the γ-region, and the pentameric assembly is referred to here as the DnaX-complex.

Interestingly, all eight maps contained peripheral, low-resolution density visible at lower display thresholds (**Figure S5**), consistent with the presence of the χψ heterodimer. Previous crystallographic and solution studies demonstrated that χψ associates flexibly with the collar domain of the DnaX-complex and can interact with single-stranded DNA–binding protein (SSB)^16,28,36,37^. However, the precise mode of χψ association with the clamp loader has remained unclear.

By applying a focused mask and extensive 3D classification, we obtained a 5.5-Å map corresponding to the χψ heterodimer and merged this density with the first clamp-loading intermediate captured here. The χψ density fits well with the previously reported crystal structure and binds the collar domain of the clamp loader via the ψ subunit. This composite map provides the first overall view of the χψ heterodimer assembled within the DnaX-holo-complex during β-clamp loading.

In the earliest loading intermediate with the 10-nt gapped DNA, the 5′-dsDNA is not resolved, suggesting that it remains flexible outside the clamp−clamp loader complex. The DNA 3′-recessed end is positioned ∼3.5 Å from Y316 but is not unwound, and the 3′-dsDNA binds to the outer surface of the open β-clamp via the basic patch, closely resembling the DNA gap recognition state with the 1-nt gapped DNA (**Figure 4C-D**).

We then reassessed the loading efficiency of β-clamp basic patch mutant using a DNA substrate containing a large gap. In contrast to the severe defects observed on 6-nt gapped DNA, the loading efficiencies of the basic patch mutants increased modestly to ∼50% of wild-type levels on this substrate. These results further underscore the importance of the β-clamp basic patch in clamp loading, particularly during the early stages of DNA recruitment (**Figure 4E**).

### 8. The intermediates depict a complete clamp-loading cycle

The eight cryo-EM 3D maps obtained with the 10-nt gapped DNA collectively describe a near-complete β-clamp loading cycle. Among these structures, six ternary clamp loader−clamp−DNA intermediates illustrate the progressive transition of the β-clamp gate from a fully open to a fully closed state as it becomes loaded onto DNA (**Figure 5, Supplemental Video 3**). These intermediates are consistent with, and complementary to, recently reported structures of the *E. coli* DnaX-complex loading β-clamp onto tailed DNA substrtes^16,17^. Overall, the observed sequence of events resembles the multistep clamp loading processes described for eukaryotic PCNA and the 9-1-1 clamp by RFC and Rad24-RFC, respectively^13–15,31^.

**Figure 5.**
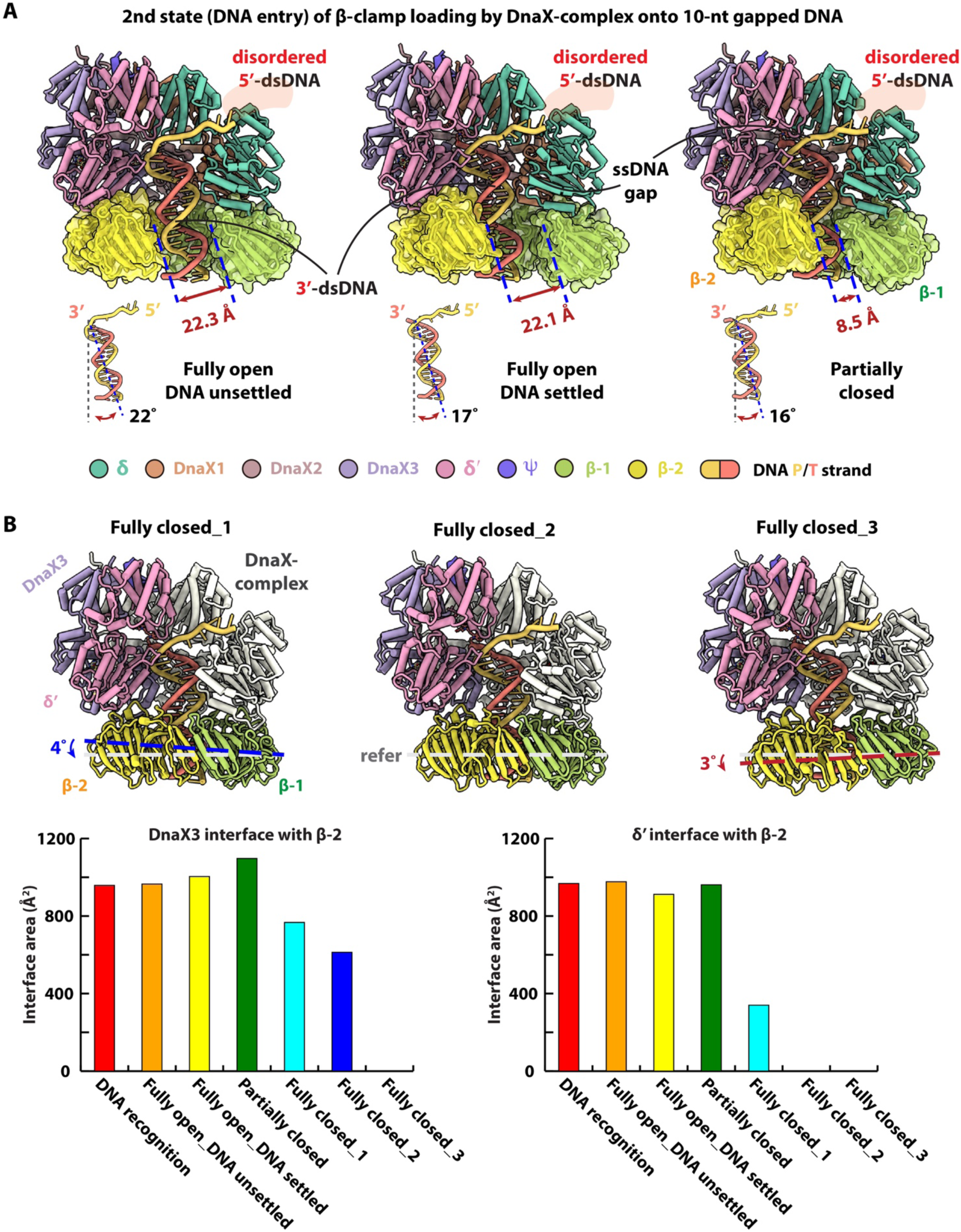
Six conformers in the second loading state of the DnaX-complex during β-clamp loading onto a 10-nt gapped DNA substrate. (**A**) In conformers 1−3, progressive settling of the 3′-dsDNA into the β-clamp is accompanied by a reduction in DNA tilt from 22° to 17° to 16°. Concurrently, the β-clamp gate narrows from 22.3 Å to 22.1 Å and then to 8.5 Å, illustrating stepwise clamp closure. (**B**) In conformers 4–6, in which the β-clamp gate is fully closed, the β-clamp progressively dissociates from the clamp loader, as indicated by clamp tilting away from the loader by ∼4° and an additional ∼3°, respectively. The lower panels show the corresponding interface areas between the DnaX3 and β2 subunits (left) and between δ′ and β2 subunits (right), highlighting changes in protein-protein interactions during clamp loader release.

Across the six loading intermediates, the clamp loader components adopt highly similar conformations, with RMSD_Cα_ values ranging from 0.5 to 0.7 Å (**Figure 5A**−**B**). In the first conformer, the β-clamp gate is open by ∼22 Å—sufficient to accommodate passage of the ∼20 Å-wide duplex DNA—and the 3′-dsDNA has entered the clamp but remains tilted by ∼22° relative to the clamp axis. We refer to this intermediate as the “fully open, DNA unsettled” state (**Figure 5A**). In the second conformer, the DNA tilt decreases to ∼17°, yielding the “fully open, DNA settled” state. In the third conformer, the clamp gate narrows further to ∼8.5 Å, and the DNA tilt decreases slightly to ∼16°.

In the fourth through sixth conformers, the clamp gate is fully closed, and the clamp progressively dissociates from the DnaX-complex. From the fourth to the fifth conformer, the clamp tilts away from the loader by ∼4°, followed by an additional ∼3° tilt from the fifth to the sixth conformer (**Figure 5B**). Quantitative analysis of protein-protein interfaces reveals that, from the DNA gap recognition state to the partially-closed clamp gate conformer, the interface area between DnaX3 and β2 increases modestly from ∼958 Å^2^ to ∼1097 Å^2^, before decreasing to ∼767 Å^2^, ∼613 Å^2^, and finally to zero in the subsequent fully closed conformers. In contrast, the interface between the δ′ subunit and β2 remains relatively constant (∼950 Å^2^) across the first four conformers, then decreases sharply to ∼340 Å^2^ in the first fully closed conformer and is completely lost in the final two conformers (**Figure 5B**).

In addition to the seven clamp loading intermediates, we captured a DnaX-complex-only structure at 2.95 Å resolution. This structure was initially presumed to represent a pre-loading state; however, careful nucleotide modeling revealed that, whereas all ATP-binding sites in the 12 clamp–clamp loader–DNA complexes (with either 1-nt or 10-nt gapped DNA) contain ATPγS, the DnaX-complex-only structure contains ADP at the ATPase sites of DnaX subunits 1 and 3, with ATPγS bound at subunit 2 (**Figure S6**). These observations indicate that ATPγS hydrolysis occurred following clamp loading and clamp loader ejection.

This finding is notable given that ATPγS is considered a slowly hydrolyzable ATP analog^12–15^. The observed hydrolysis suggests that the *E. coli* DnaX-complex is a more robust ATPase than its eukaryotic counterpart, consistent with our biochemical results, showing that, although DnaX-complex does not unwind DNA at nicks or very small gaps, it can nevertheless load the β-clamp onto such substrates given sufficient time (**Figure 3A**).

### 9. The DNA recessed 5′ end does not stimulate clamp loading activity of DnaX-complex

In the absence of DNA, clamp loaders from *E. coli*, yeast and humans adopt broadly similar overall architectures (**Figure S7A**). Upon DNA binding, however, the eukaryotic RFC clamp loader undergoes conformational changes that expose basic residues on the Rfc1 AAA+ module, together with the unique Rfc1 BRCT domain, forming a large, highly positively charged surface at the shoulder. This surface serves as a secondary binding site that stably engages the 5′-dsDNA segment during clamp loading^12,17,31,35^. In contrast, with either 1-nt or 10-nt gapped DNA, the 5′-dsDNA remains largely disordered at the shoulder region of the bacterial DnaX-complex, as described above.

Electrostatic surface potential analysis further reveals that the shoulder region of the DnaX-complex is largely neutral or weakly negatively charged, in stark contrast to the highly positively charged shoulder of eukaryotic RFC (**Figures 6A and S7B-D**). This difference suggests that the bacterial clamp loader lacks a dedicated shoulder DNA-binding site capable of stably engaging the 5′-dsDNA.

**Figure 6.**
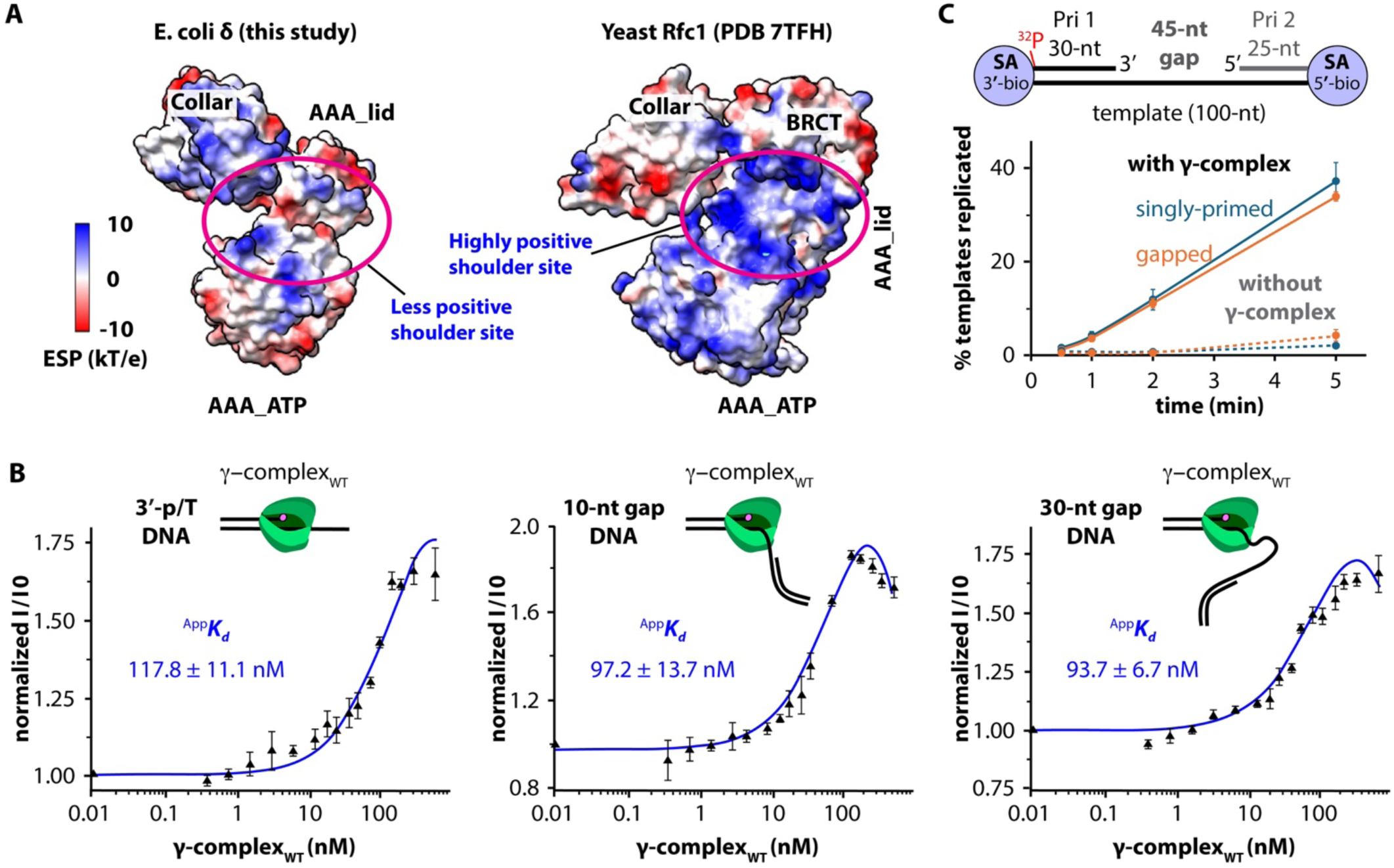
The DnaX-complex shoulder is not highly positively charged and is not required for efficient clamp loading. (**A**) Comparison of electrostatic surface potentials showing that the eukaryotic clamp loader subunit Rfc1 (PDB: 7TFH) contains a highly positively charged dsDNA binding site at the shoulder region (purple oval), whereas the corresponding region of *E. coli* δ (this study) lacks a comparably charged surface. (**B**) Fluorescence anisotropy measurements indicate that the presence of a 5′-dsDNA does not significantly enhance DNA binding by the DnaX-complex. DNA substrates consisting of a tailed primer/template with only a recessed 3′ end (left), a 10-nt gapped DNA (middle), or a 30-nt gapped DNA (right) bind the DnaX-complex with similar apparent affinities (^App^*K_d_* values of 118 nM, 97 nM, and 94 nM, respectively). Fluorescence signals were normalized (I /I0) to the signal obtained in the absence of protein. Assays were performed in triplicate, and ^App^*K_d_* values are shown as mean values (black triangles) ± standard deviations (error bars). (**C**) The rate of β-clamp loading onto a DNA containing only a recessed 3′ end is comparable to that observed for loading onto a 45-nt gapped DNA containing both recessed 3′ and 5′ ends.

If the DnaX-complex does not possess a functional shoulder DNA-binding site, then the presence of a 5′-recessed end should not enhance clamp loader-DNA interactions or clamp loading efficiency. To test this prediction, we measured the affinity of the γ-complex for three different DNA substrates using a fluorescence polarization-based binding assay with Cy3-labeled DNA (see **STAR METHODS**): (1) a primer/template (p/T) DNA containing only a recessed 3′ end, (2) a 10-nt gapped DNA containig both recessed 3′ and 5′ ends, and (3) a 30-nt gapped DNA containing both recessed ends (**Figure 6B**). The apparent dissociation constant (^App^*K_d_*) for these substrates were 117.8 ± 11.1 nM, 97.2 ± 13.7 nM, and 93.7 ± 6.7 nM, respectively, indicating that the presence of a recessed 5′ end does not significantly enhance DNA binding by the DnaX-complex.

We examined whether the presence of a 5′-recessed end influences clamp loading activity using a clamp loading-dependent primer extension assay. In this assay, DNA synthesis by Pol III requires loading of the β-clamp and thus serves as a proxy for clamp loading efficiency (see **STAR Methods**). We used two DNA substrates. The first was a p/T DNA, consisting of a recessed 5′ end ^32^P-labeled 30-nt primer (Pri-1) annealed to a 100-nt template biotin-labeled at both ends, thereby containing only a 3′-recessed end. The second was a 45-nt gapped DNA, generated by annealing an additional 25-nt primer (Pri-2), thereby creating both 3′- and 5′-recessed ends. Streptavidin (SA) was added to block the DNA ends and prevent the β-clamp from diffusing off the DNA (**Figures 6C and S8A**). Sufficient Pol III was added to ensure that the reaction was rate-limited only by clamp loading. The p/T and gapped DNAs were not extended by Pol III in the absence of clamp loading under our conditions and thus serve as negative controls. Both substrates were extended in the presence of clamp loading. We found that the amount of synthesized DNA was similar for both substrates, with the p/T DNA yielding a slightly higher amount (**Figure 6C**). This result is again consistent with the structural observations.

We further examined stimulation of DnaX-complex ATPase by DNA substrates containing a nick or gaps of different sizes in the presence of β-clamp (**Figure S8B**). Small and larger gapped DNAs produced similar levels of ATPase hydrolysis, whereas a 5-nt gapped DNA yielded a measurably lower ATPase activity. Considering that clamp loading efficiency onto nicked or 2-nt gapped DNA is markedly lower than onto DNA with 4-nt or larger gap (**Figure 3A**), these results imply that a significant fraction of ATP hydrolysis might be attributed to failed DNA bending events during clamp loading, even though these substrates consume a similar amount of ATP as the 10-nt gapped DNA. By contrast, the reduced ATP consumption observed with the 5-nt gapped DNA may indicate that its 5′-dsDNA segment can more readily bend back to engage the β-clamp basic patch (as observed in the state 2 structures captured with 1-nt gapped DNA), thereby facilitating clamp loading, specifically the entry of the 3′-dsDNA into clamp–clamp loader chamber.

## DISCUSSION

In this work, we uncover a fundamental mechanistic difference between the bacterial and eukaryotic clamp loading onto nicked and small-gapped DNA substrates. A minimum stretch of ∼ 6 nt of ssDNA is generally required to thread through the clamp loader chamber to enable recognition and binding of a recessed DNA 3′ end for clamp loading. When the gap is smaller than this threshold, eukaryotic RFC widens the gap by unwinding the DNA from the 3′ end, whereas the bacterial DnaX-complex instead sharply bends the 5′-dsDNA by ∼150°, forcing the 3′-dsDNA into the clamp chamber without detectable strand separation. Despite the absence of DNA unwinding activity, the δ-subunit residue Y316 of the DnaX-complex approaches and interacts with the recessed 3′ end, consistent with a role in DNA end recognition rather than strand separation.

Importantly, these mechanistic distinctions are based on the substrates and conditions examined in this study. While our data show no evidence of DNA unwinding by the bacterial clamp loader under these conditions, we cannot exclude the possibility that DNA bending also occurs in eukaryotic systems or that DNA unwinding may contribute to bacterial clamp loading under alternative physiological conditions, in the presence of additional factors, or with different DNA substrates.

Small ssDNA gaps and nicks are common intermediates generated during DNA damage processing and repair. The ability of the DnaX-complex to load the β-clamp efficiently at such sites—without requiring DNA unwinding or a dedicated shoulder DNA-binding site—suggests a mechanism that may be particularly well suited for coordinating DNA repair synthesis, and would be consistent with repair deficiency in yeast when using RFC that cannot bind a small gap due to loss of its 5′-dsDNA binding domain (N-terminal truncation)^14,38^. Consistent with this idea, the *E. coli* β-clamp is known to play central roles in recruiting and coordinating multiple DNA repair and translesion synthesis polymerases, as well as other repair-associated factors^39–49^. Sharp DNA bending at small gaps may therefore facilitate rapid clamp loading and polymerase exchange at sites of damage, enabling efficient downstream repair.

### Proposed β-clamp loading mechanism by DnaX-complex

The bacterial clamp loader appears to load the β-clamp in a DNA structure–dependent manner. During normal bacterial replication, Okazaki fragments are typically 1−2 kb long, such that the preceding Okazaki fragment (i.e., 5′-dsDNA) is too distant to influence clamp loading and can effectively be ignored. Under these conditions, the 3′-dsDNA first binds in a tilted orientation to the basic patch on the outer surface of the open β-clamp (**Figure 7**). The DnaX-complex then engages the recessed primer 3′ end via the δ-subunit Υ316, allowing the 3′-dsDNA to diffuse toward the higher-affinity binding site within the β-clamp chamber through a series of steps captured here and in recent studies^16,17^.

**Figure 7.**
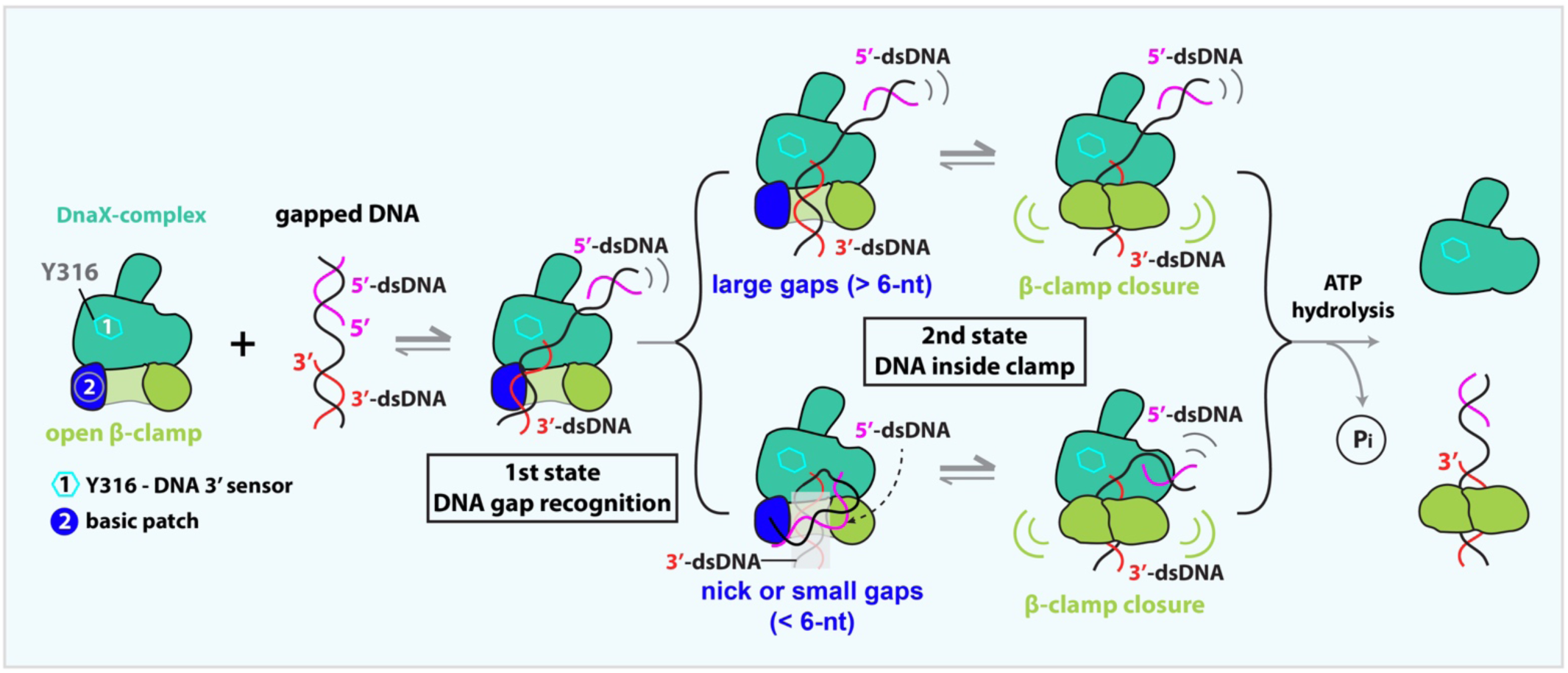
Proposed DNA structure–dependent mechanism for β-clamp loading by the DnaX-complex. The bacterial clamp loader recognizes, but does not unwind, the recessed DNA 3′ end via the δ-subunit Y316 residue, and the β-clamp contains a conserved basic patch on its outer surface that can bind dsDNA. When a gapped DNA substrate encounters a DnaX-complex bound to an open β-clamp, the 3′-dsDNA binds the β-clamp basic patch in a tilted orientation (first state). If the ssDNA gap is sufficiently long to thread through the clamp loader (>6 nt), the 5′-dsDNA remains outside the loader while the 3′-dsDNA diffuses to the higher affinity interior chamber of the β-clamp (second state). In contrast, at a nick or a small ssDNA gap (<6 nt), the adjacent 5′-dsDNA bends sharply toward the 3′-dsDNA, adopts a U-shaped conformation that promotes displacement of the 3′-dsDNA toward the β-clamp chamber. Once the 3′-dsDNA enters the clamp interior, electrostatic interactions between the negatively charged DNA backbone and the positively charged inner surface of the β-clamp are proposed to drive clamp gate closure. Subsequent ATP hydrolysis triggers dissociation of the DnaX-complex, leaving the β-clamp encircling the 3′-dsDNA.

When the DnaX-complex encounters a nick or a small gap (<6 nt), the loading pathway diverges. In contrast to eukaryotic RFC, which can unwind the DNA 3′ end by up to 6 bp using the Rfc1 separation pin W638—a two-ring aromatic residue—while the corresponding residue in *E. coli*, δ-subunit Y316, is a single-ring aromatic residue and does not unwind DNA^14^. Instead, the bacterial clamp loader appears to have evolved a distinct strategy that may be particularly suited for recruiting repair polymerases (Pol I, II, IV, and V) to sites of DNA damage^47,50–53^.

At small gaps, the primer 3′ end is initially sterically hindered from entering the DnaX-complex chamber because the adjacent 5′-dsDNA segment is too bulky to be accommodated. In the absence of stable 5′-dsDNA binding at the clamp loader shoulder, the 5′-dsDNA bends sharply down toward the 3′-dsDNA, generating a U-shaped DNA conformation (**Figure 7**). This sharp bending has two important consequences. First, it exposes the recessed primer 3′ end, facilitating its recognition by δ Υ316 and its subsequent entry and proper positioning within the clamp loader chamber (**Figure S4**). Second, the bending-induced strain within the DNA promotes the displacement of the 3′-dsDNA from the external β-clamp binding site and drives its insertion into the β-clamp chamber.

Following entry of the 3′-dsDNA into the clamp, the 5′-dsDNA binds transiently to the β-clamp basic patch vacated by the 3′-dsDNA, further promoting clamp closure. Electrostatic attraction between the negatively charged DNA backbone and the positively charged interior surface of the β-clamp is expected to drive full or near-complete gate closure, analogous to mechanisms proposed for PCNA and the 9-1-1 clamp^12–15,18,25–27,31^. Clamp closure is then coupled to ATP hydrolysis, triggering dissociation of the DnaX-complex. Upon clamp loader release, the bent DNA is expected to relax back to a linear configuration, allowing the β-clamp to fully encircle the DNA. Such a mechanism is likely advantageous at discontinuous DNA substrates encountered during DNA repair, where rapid clamp loading and polymerase exchange are required. Consistent with this model, our loading assays demonstrate that the β-clamp can be successfully loaded onto a 1-nt gapped DNA substrate (**Figure 3A**).

### The β-clamp external DNA binding site may compensate for the lack of a shoulder 5′ DNA site in DnaX-complex

Sharp DNA bending generally requires at least two spatially separate DNA-binding sites. For example, during replication origin recognition, yeast origin DNA is bent by ∼50° by the origin recognition complex (ORC), which engages DNA at two distinct sites—one within the ORC ring and the other on the external surface formed by Orc5 and Orc6^54,55^. Similarly, eukaryotic RFC employs two dsDNA-binding sites to load PCNA onto gapped DNA: the primary site within the chamber that binds the 3′-dsDNA, and a secondary, highly positively charged shoulder site that engages the 5′-dsDNA^13,15,25–27^.

In contrast, the *E. coli* DnaX-complex lacks a comparable positively charged shoulder region, and the 5′-dsDNA does not stably bind there. Instead, our structures and biochemical data indicate that the β-clamp itself has evolved a conserved basic patch on its external surface that can bind DNA, providing the second DNA-binding site for the bacterial clamp loading system (**Figures 2D-E** and **S7A-C**). This β-clamp basic patch may therefore functionally compensate for the absence of a shoulder 5′-dsDNA-binding site in the DnaX-complex.

Notably, the RFC shoulder site includes the Rfc1 N-terminal BRCT domain, which is unique to eukaryotes and has been proposed to function in initial DNA recruitment^3,15,25–27^. By analogy, the β-clamp basic patch may serve a related role in bacterial clamp loading by stabilizing transient DNA interactions during recruitment and loading. Thus, although the molecular components differ, both systems appear to utilize two DNA-binding sites to achieve clamp loading at nicks and gaps, highlighting a convergent functional strategy implemented through distinct structural solutions.

### DnaX-complex is anchored to the replication fork for repeated clamp loading

The DnaX-complex functions as more than a clamp loader; it also serves as a central organizer of the bacterial replisome, coordinating helicase and polymerase activities at the replication fork^56–59^ (**Figure S9**). This multifunctionality is conferred by the CTDs of the three τ subunits (τ_CTDs_), which mediate interactions with multiple replication proteins^60^. One τ_CTD_ anchors the leading strand Pol III, a second τ_CTD_ anchors the lagging-strand Pol III, and the third can recruit an additional polymerase, providing backup or facilitating completion of lagging strand synthesis when polymerase exchange is required^61,62^.

In addition to polymerase tethering, the DnaX-complex is linked to the replication fork through its χ subunit, which binds the SSB^36,57^. This interaction helps position the clamp loader at sites of active DNA synthesis and likely promotes efficient and repeated β-clamp loading at newly generated primer-template junctions. In this study, we visualized for the first time the χψ heterodimer assembled within the DnaX-holo-complex during β-clamp loading (**Figure S9**), providing structural context for earlier biochemical and solution studies^36,57,63^.

The ability of the DnaX-complex to remain anchored at the replication fork offers a mechanistic explanation for how clamp loading can be efficiently coupled to ongoing DNA synthesis. By maintaining proximity to a primer-template junction and to SSB-coated ssDNA, the DnaX-complex is well positioned to rapidly respond to newly formed primers during replication and repair, ensuring timely loading of the β-clamp as needed.

Beyond its biological implications, the distinct clamp-loading mechanism described here may also have biomedical relevance. The reliance of bacterial clamp loading on a conserved β-clamp DNA-binding patch and on sharp DNA bending represents features that are absent from eukaryotic clamp loaders. These mechanistic differences raise the possibility that bacterial clamp loading could be selectively targeted, providing a potential avenue for the development of species-specific antibacterial strategies that interfere with DNA replication and repair without affecting eukaryotic counterparts.

### Limitations of the Study

This study reveals a distinct mechanism by which the *E. coli* DnaX-complex loads the β-clamp onto nicked and small-gapped DNA substrates (<6 nt), characterized by sharp DNA bending rather than DNA unwinding, as observed for eukaryotic RFC. Although these features are well supported by in vitro cryo-EM and biochemical analyses, the biological consequences of these mechanistic differences in vivo remain to be fully established. Moreover, we cannot exclude the possibility that DNA bending also occurs in eukaryotic systems, or that DNA unwinding may contribute to bacterial clamp loading under physiological conditions or with DNA substrates not examined here.

In particular, the physiological roles of the conserved basic patch on the β-clamp and of the δ-subunit residue Y316 during DNA replication and repair warrant further investigation. While our structural and biochemical data indicate that these elements are important for DNA recognition and clamp loading at small gaps, their precise contributions to cellular DNA repair pathways remain unknown. Determining how these features influence polymerase selection, clamp dynamics, and repair efficiency in vivo will be an important direction for future studies.

In addition, although our experiments capture multiple clamp-loading intermediates under defined in vitro conditions, it remains possible that alternative DNA substrates, accessory factors, or cellular environments could modulate the clamp loading pathway. Thus, while our results establish a mechanistic framework for understanding bacterial clamp loading at nicks and small gaps, further work will be required to define how this mechanism operates within the full cellular replication and repair machinery.

## STAR ★ METHODS

Detailed methods are provided in the online version of this paper and include the following:

- KEY RESOURCES TABLE
- RESOURCE AVAILABILITY

- Lead contact
- Materials availability
- Data and code availability
- EXPERIMENTAL MODEL AND SUBJECT DETAILS
- METHOD DETAILS

- Protein expression and purification
- β-clamp mutants preparation
- γ-complex mutant preparation
- Gel filtration-based analysis of β-clamp loading
- Fluorescence polarization-based DNA binding assay
- Rate of clamp loading onto primed versus gapped DNA
- Cryo-EM grids preparation and data collection
- Image processing and 3D reconstruction
- Model building, refinement, and validation
- QUANTIFICATION AND STATISTICAL ANALYSIS

## SUPPLEMENTAL INFORMATION

Supplemental Information can be found online at:

## ACKNOWLEDGMENTS

Cryo-EM micrographs were collected at the David Van Andel Advanced Cryo-Electron Microscopy Suite in the Van Andel Institute. We thank G. Zhao and X. Meng for assistance with data collection. This work was supported by U.S. National Institutes of Health grants GM115809 (to M.E.O.) and GM131754 (to H.L.), the Howard Hughes Medical Institute (to M.E.O.), and the Van Andel Institute (to H.L.).

## AUTHOR CONTRIBUTIONS

F.Z., M.E.O., and H.L. designed research; F.Z., N.Y., R.E.G. and M.L. performed research; F.Z., R.E.G., N.Y.Y., M.L., M.E.O., and H.L. analyzed the data; F.Z., M.E.O., and H.L. wrote the manuscript with input from all authors.

## DECLARATION OF INTERESTS

The authors declare no competing interests.

## STAR ★ METHODS

### KEY RESOURCES TABLE

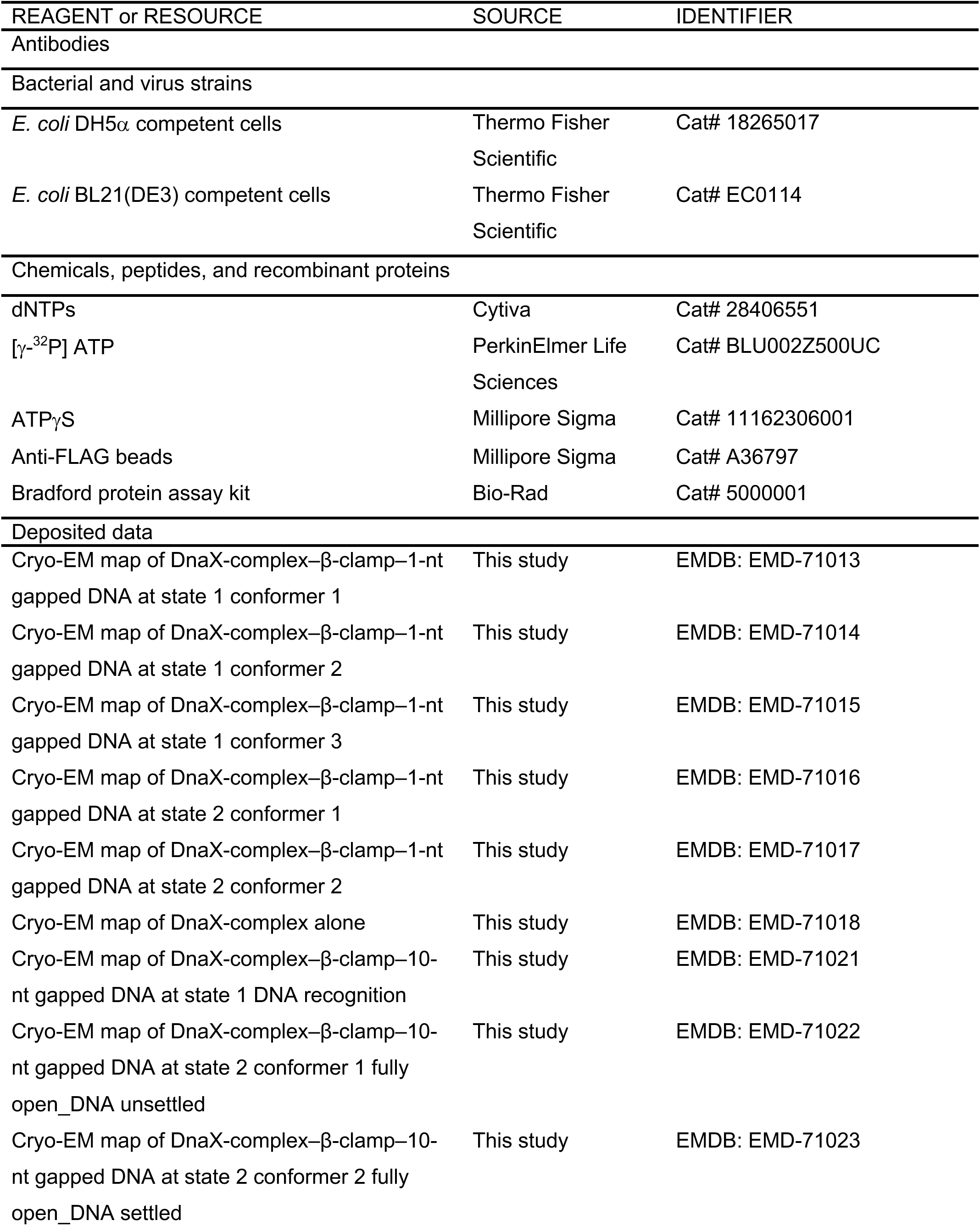
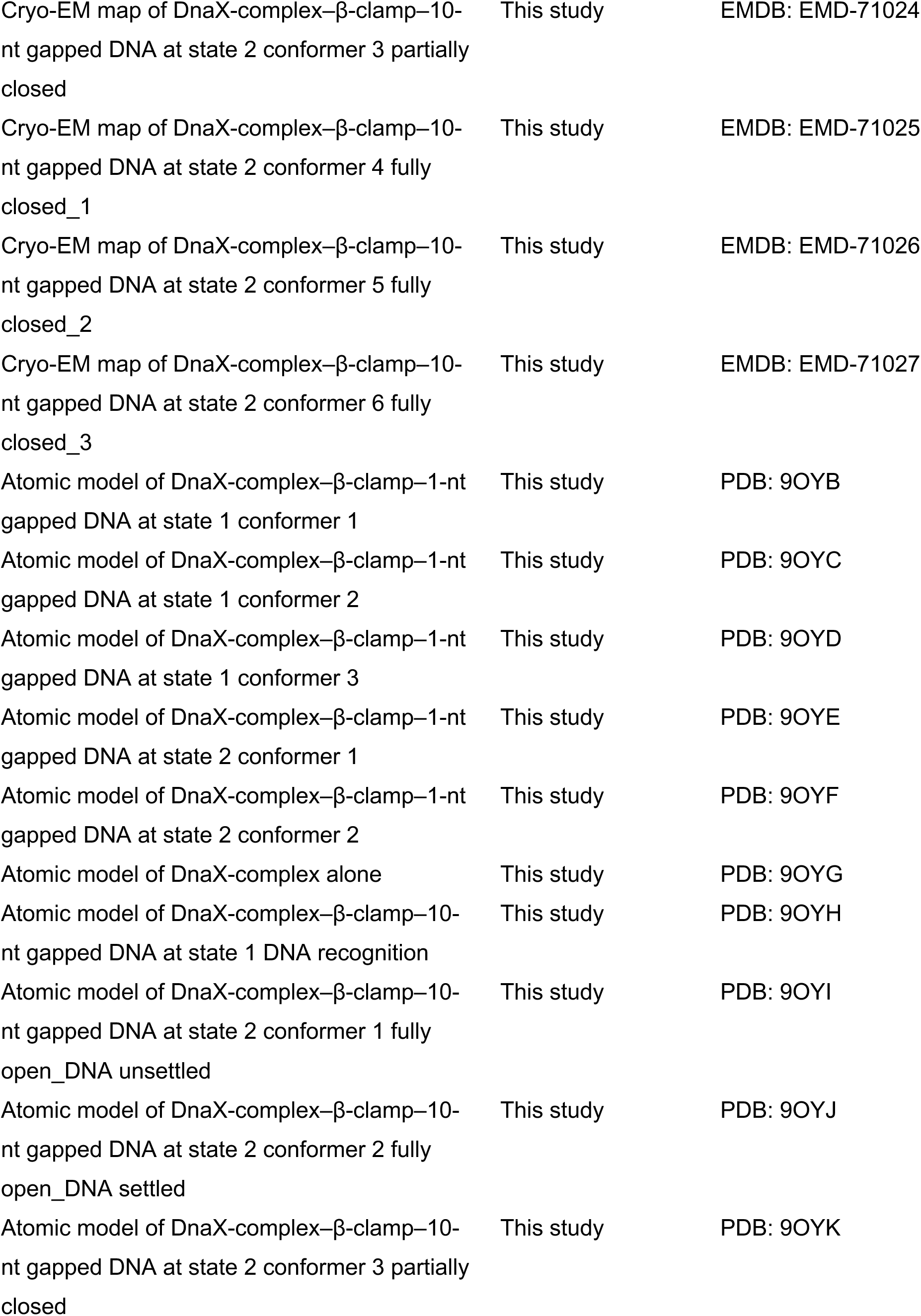
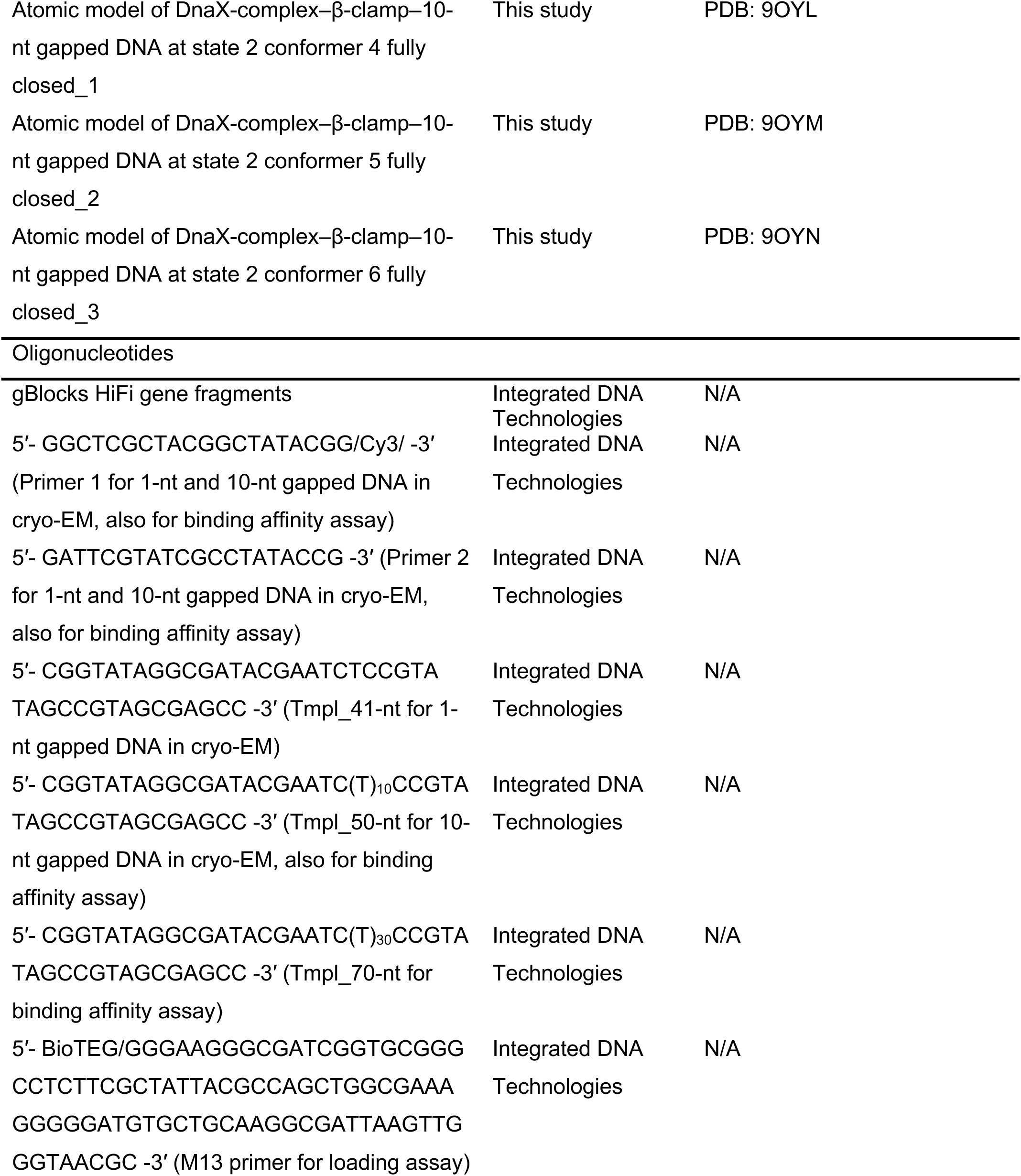
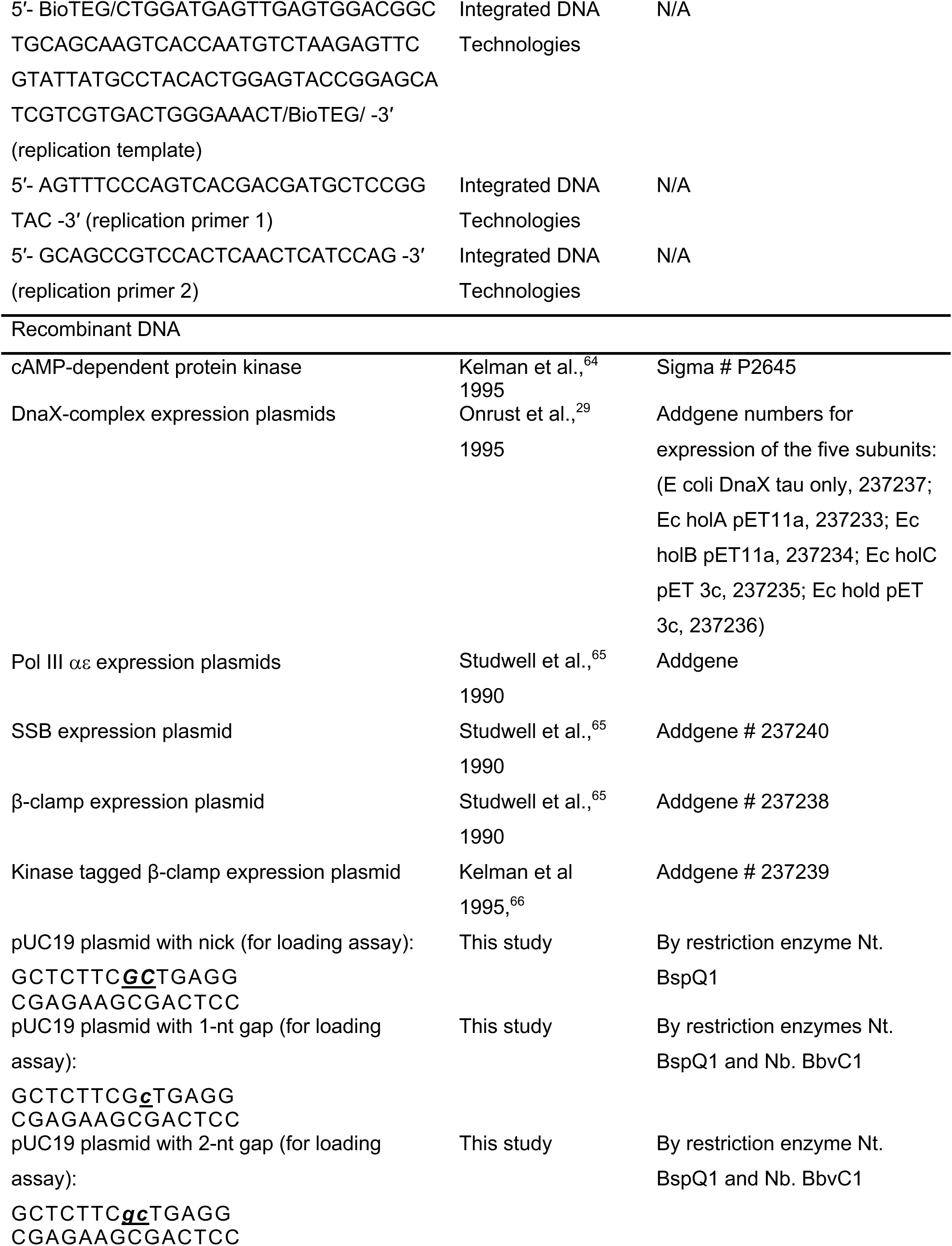
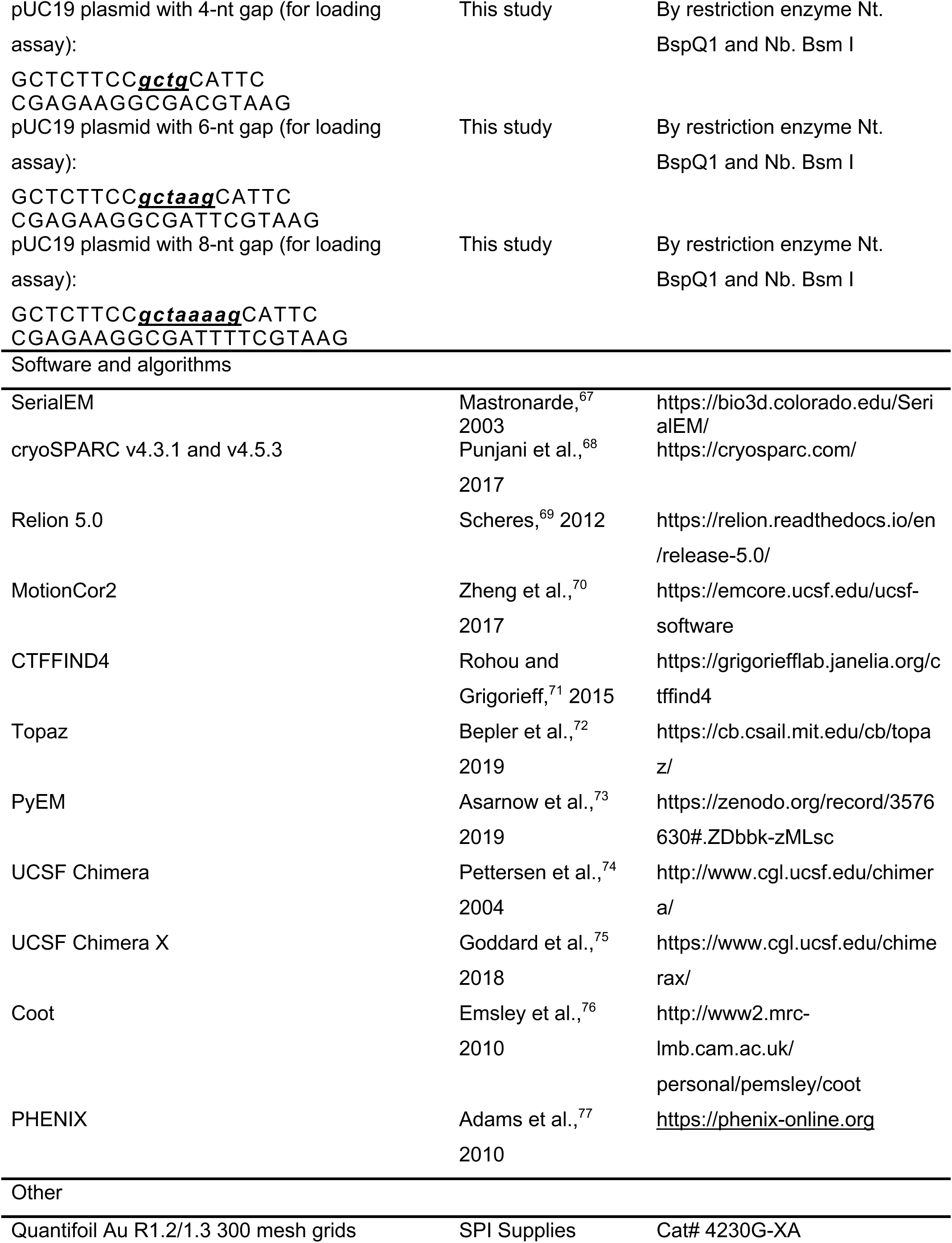
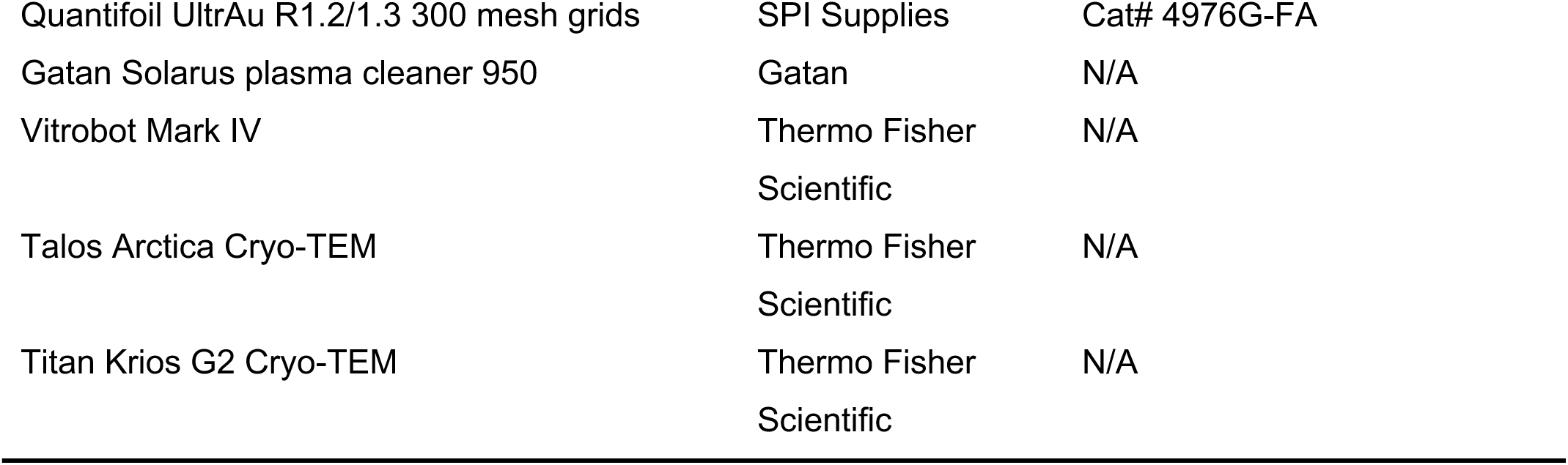

## RESOURCE AVAILABILITY

### Lead contact

Further information and requests for reagents and resources should be directed to, and will be fulfilled by, the Lead Contact, Huilin Li.

### Materials availability

All plasmid clones used to express the proteins in this study are available upon request without restrictions, although a material transfer agreement (MTA) may be required.

### Data and code availability

- Five conformers of the DnaX-complex bound to β-clamp and 1-nt gapped DNA (states 1 and 2) have been deposited in the Electron Microscopy Data Bank under accession codes EMD-71013, EMD-71014, EMD-71015, EMD-71016, and EMD-71017. The corresponding atomic models have been deposited in the Protein Data Bank under accession codes 9OYB, 9OYC, 9OYD, 9OYE, and 9OYF. These data will be publicly available upon publication.
- One DnaX-complex–alone structure and seven conformers of DnaX-complex bound to β-clamp and 10-nt gapped DNA have been deposited in the Electron Microscopy Data Bank under accession codes EMD-71018, EMD-71021, EMD-71022, EMD-71023, EMD-71024, EMD-71025, EMD-71026 and EMD-71027. The corresponding atomic models have been deposited in the Protein Data Bank under accession codes 9OYG, 9OYH, 9OYI, 9OYJ, 9OYK, 9OYL, 9OYM, and 9OYN. These data will be publicly available upon publication.
- This paper does not report original code.
- Clones used for expression of factors in this study have been deposited in the Addgene repository.
- Any additional information required to reanalyze the data reported in this work will be available from the Lead Contact upon request.

## EXPERIMENTAL MODEL AND STUDY PARTICIPANT DETAILS

### Bacterial strains

Expression plasmids encoding the *E. coli* DnaX-complexes—specifically the γ-complex (δγ₃δ′χψ), used for biochemical studies, and the τ-complex (δτ₃δ′χψ), used for cryo-EM studies—as well as the β-clamp and its related mutants, SSB, and Pol III αε complex, were generated as described previously^29,65^. All proteins were expressed from the plasmids listed in the **KEY RESOURCES TABLE** following transformation into *E. coli* BL21(DE3) cells (Thermo Fisher Scientific) and cultured in LB media.

## METHOD DETAILS

### Protein expression and purification

The *E. coli* DnaX-complex subunits δ, γ, τ, δ′, ψ, and χ were purified as described previously^29^. The β-clamp, Pol III αε complex, and SSB were purified as described previously^65^. The clamp loader γ-complex was reconstituted from purified subunits and further purified by ion-exchange chromatography using a Cytiva MonoQ column as described^29^. Briefly, purified subunits (1–4 mg each in approximately 3 mL of 20 mM Tris-HCl, pH 7.5, 1 mM DTT, 0.5 mM EDTA, 10% glycerol) were mixed at a molar ratio of δ:γ₃:δ′:ψ:χ = 1.5:1:1.5:1.5:2. The subunit mixture was incubated for 30 min at 15 °C and then applied to a 1-mL MonoQ column (GE Healthcare, Chicago, IL). Proteins were eluted using a 10-column-volume linear gradient of Buffer A (20 mM Tris-HCl, pH 7.5, 0.5 mM DTT, 1 mM EDTA, 10% glycerol) supplemented with 50 mM NaCl, and 0.2-mL fractions were collected. Fractions were analyzed by 12% SDS–PAGE. The γ-complex eluted at approximately 0.4 M NaCl, whereas individual subunits and subcomplexes eluted earlier. The τ-complex was reconstituted and purified in the same procedure, except that τ was substituted for γ. Peak fractions of each complex were pooled and dialyzed against Buffer A supplemented with 50 mM NaCl, aliquoted, snap-frozen in liquid nitrogen, and stored at −80 °C.

### Mutant β-clamp preparation

gBlocks HiFi gene fragments were designed to contain the full-length *dnaN* gene, including an N-terminal 3×FLAG tag followed by a protein kinase A recognition site (LRRASLG)^66^. The gBlocks were cloned into pRSFDuet (Novagen, Burlington, MA) that had been digested with NcoI-HF and BamHI-HF and treated with alkaline phosphatase. This strategy was used to generate wild-type β-clamp (β-clamp_WT_) as well as two mutants containing three amino acid substitutions: R96A, R103A, and R105A (β-clamp_Mut1_) and R96E, R103E, and R105E (β-clamp_Mut2_).

The gBlocks were dissolved in 20 mM Tris-acetate (pH 7.9), 50 mM K-acetate, 10 mM MgCl_2_, and 100 µg/mL BSA, followed by digestion with NcoI-HF and BamHI-HF. The resulting plasmids were transformed into *E. coli* DH5α cells (ThermoFisher Scientific, Waltham, MA), plated on LB agar containing kanamycin (50 µg/mL), and incubated overnight at 37 °C. Colonies were screened by miniprep and NcoI-HF/BamHI-HF restriction digestion on native agarose gels, followed by maxiprep and verification of the entire *dnaN* coding sequence by Sanger sequencing (Genewiz, South Plainfield, NJ).

Sequence-verified plasmids were transformed into *E. coli* BL21(DE3) cells, and 3 L cultures were grown to an OD_600_ of 0.6 before induction with 2 mM IPTG for 12 h at 15 °C. Cells were harvested by centrifugation, resuspended in Buffer A supplemented with 750 mM NaCl, and lysed using an Avestin EmulsiFlex-C-50 (Ottawa, ON, Canada). PMSF was added to a final concentration of 1 mM, and the lysate was clarified by centrifugation at 12,500 rpm in an F14-6×250 rotor (ThermoFisher Scientific, Waltham, MA) at 4 °C for 1 h.

Anti-FLAG affinity purification was performed by incubating the clarified supernatant (∼400 mL) with 5 mL anti-FLAG resin for 30 min at 4 °C with gentle mixing. The beads were collected by centrifugation and washed once with 450 mL Buffer A containing 300 mM NaCl, followed by three additional wash with 400 mL Buffer A containing 300 mM NaCl. The final bead pellet was resuspended in 15 mL Buffer A containing 50 mM NaCl and loaded onto an XK-16/20 column (Cytiva, Marlborough, MA). The column was washed with 15 mL Buffer A containing 1 M NaCl and then equilibrated with Buffer A containing 50 mM NaCl. Bound proteins were eluted in four successive 30-min pulses of 5 mL Buffer A containing 50 mM NaCl and 0.23 mg/mL 3×FLAG peptide. Eluted fractions were analyzed by SDS–PAGE, pooled, dialyzed against Buffer A, concentrated to approximately 2 mg/mL, and stored at −80 °C.

### Mass photometry

Data were collected using a OneMP mass photometer (Refeyn), calibrated with bovine serum albumin (66 kDa), β-amylase (224 kDa), and thyroglobulin (670 kDa). Movies were acquired for 6,000 frames (60 s) using AcquireMP software (version 2.4.0) with default settings. Final protein concentrations were empirically adjusted to achieve approximately 75 binding events per second. Peak mass values are expected to fall within ∼5% error. Raw data were converted to frequency distributions using Prism 9 (GraphPad) and a bin size of 10 Da.

### Mutant γ subunit preparation

The δ-subunit Y316A mutant was generated using a HiFi DNA fragment encompassing the region of *holA* (gene encoding δ) from the internal AflII site to the BamHI site downstream of the stop codon, including placement of a C-terminal 3×FLAG tag. This gBlock was inserted into AflII/BamHI-digested pET11-holA to generate pET11-holA(Y316A). The full sequence of the mutant *holA* gene was verified by Sanger sequencing.

The pET11-holA(Y316A) plasmid was transformed into *E. coli* Bl21(DE3) cells, and 12 L cultures were grown under ampicillin selection, induced with 2 mM IPTG, and lysed in an Avestin EmulsiFlex-C-50 (Ottawa, ON, Canada). The δ Y316A protein was purified as described above for the β-clamp mutants, concentrated, and stored at −80 °C.

The γ-complex containing the δ Y316A mutation was reconstituted by mixing purified δ Y316A mutant with γ, δ‘, χ, and ψ subunits and purified as described previously^63^ and as briefly outlined above.

### Construction of DNA substrates for the β-clamp loading assay

pUC19 plasmid DNA was engineered to generate nicked or gapped DNA substrates of defined gap sizes using one or two nicking endonucleases. The required insertions into pUC19 are listed in the **KEY RESOURCES TABLE**. The nicking enzymes cut the same DNA strand, producing a short single-stranded region that can be released from the circular plasmid to generate a nick or gap of defined length.

For generation of 1- and 2-nt gapped DNA substrates, Nb.BbvCI and Nt.BspQI were used. For generation of 4-, 6-, and 8-nt gapped DNA substrates, Nb.BsmI and Nt.BspQI were used. To generate each gapped circular DNA, 43 µg of plasmid DNA was digested in a reaction containing 50 mM Tris-HCl (pH 7.9), 100 mM NaCl, 10 mM MgCl_2_, and 100 µg/mL BSA with 40 U each of the nicking enzymes. For 1- and 2-nt gapped DNA, Nb.BbvCI and Nt.BspQI digestions were performed sequentially at 37 °C and 50 °C, respectively. For 4-, 6- and 8-nt gapped DNA, Nt.BspQI and Nb.BsmI digestions were performed sequentially at 50 °C and 65 °C, respectively.

Following digestion, enzymes were heat-inactivated at 80 °C for 30 min, and reactions were terminated by addition of EDTA to a final concentration of 5 mM. For 4-, 6- and 8-nt gapped DNA substrates, a complementary oligo trap was added (4-nt gap: 5’-TTCTTAGCTT-3’; 6-nt gap: 5’-TTCTTAGCTT-3’; 8-nt gap: 5’-TCTTTTAGCT-3’), and the reactions were heated to 55 °C and then slowly cooled to room temperature.

Approximately 25 µg of each reaction was purified using a spin column, and DNA was eluted in three successive 20-µL elutions with 5mM Tris-HCl (pH 8.5) at 70 °C. For the singly primed DNA substrate, a biotinylated M13 primer was annealed to M13 ssDNA, followed by addition of a 5-fold molar excess of streptavidin (**KEY RESOURCES TABLE**).

### Gel filtration–based β-clamp loading assay

β-Clamp loading onto DNA was analyzed using the γ-complex, isotopically ^32^P-labelled β-clamp, and the corresponding clamp and clamp loader mutants. DNA-bound ^32^P–β-clamp was separated from unbound ^32^P–β-clamp by gel filtration through a 5-mL Bio-Gel A15m column, essentially as described previously^30^. Briefly, β-clamp was radiolabeled by incubating 1 nmol FLAG–kinase–tagged wild-type or mutant β-clamp in 100 μL buffer A containing 16.7 pmol γ-^32^P-ATP and 0.45 μg protein kinase A (PKA; Sigma, St Louis, MO) for 30 min at 37°C. Radiolabeled ^32^P–β-clamp was separated from unincorporated ^32^P-ATP using a Cytiva G25 MicroSpin column equilibrated with buffer A supplemented with 50 mM NaCl. The concentration and specific radioactivity of the purified ^32^P–β-clamps were determined by Bradford assay and liquid scintillation counting.

Clamp loading reactions contained 1 pmol (10 nM) pUC19 plasmid DNA with a nick or defined gap size, or singly primed M13 ssDNA; 2 pmol (20 nM) ^32^P–FLAG–PK–β-clamp (dimer); and varying concentrations of γ-complex or their mutants, β-clamp_Mut1_, β-clamp_Mut2_ and γ-complex_Mut_ in a total volume of 100 μL Buffer B (20 mM Tris-HCl, pH7.5, 0.1 mM EDTA, 40 μg/mL BSA, 4% glycerol, 5 mM dithiothreitol) supplemented with 8 mM MgCl_2_, 2 mM ATP and 50 mM NaCl. Reactions were incubated for 5 min at 37 °C. DNA-bound ^32^P–β-clamp was separated from free ^32^P–β-clamp by gel filtration through a 5-mL Bio-Gel A15m column equilibrated with Buffer B supplemented with 8 mM MgCl_2_ and 100 mM NaCl. Fractions (180 μL) were collected, and 150 μL aliquots were quantified by liquid scintillation counting. Radioactivity was converted to femtomoles using the empirically determined specific radioactivity of ^32^P–β-clamp.

### Cryo-EM grids preparation and data collection

The 1-nt and 10-nt gapped DNA substrates used for cryo-EM studies are shown in **Figures 1C** and **4A**. *In vitro* assembly of clamp loading complexes containing purified *E. coli* DnaX-complex, β-clamp, and DNA substrates followed our previously established reconstitution procedure for the yeast Polδ–PCNA–DNA complex^78^. Briefly, either 13.0 μL of 1-nt gapped DNA at 7.1 μM or 8.0 μL of 10-nt gapped DNA at 11.5 μM were mixed with 1.0 μL β-clamp at 77.6 μM and incubated at 30 °C for 10 min. The DNA-clamp mixtures, together with 1.0 μL of 10 mM ATPγS and 1.0 μL of 100 mM Mg-acetate, were then added to 7.1 μL DnaX-complex at a concentration of 8.4 μM. Ultrapure H_2_O was added to bring the final reaction volume to 20 μL. The mixture was incubated in an ice–water bath for an additional 10 min prior to cryo-grid preparation.

Quantifoil Au or UltrAu R1.2/1.3 300-mesh grids were glow discharged for 0.5 min or 3 min, respectively, using a Gatan Solarus plasma cleaner. Aliquots (3 μL) of the assembled clamp loading mixtures were applied onto the grids, which were vitrification using a Vitrobot Mark IV (Thermo Fisher Scientific) under the following conditions: blot time of 2 s for Au grids or 2.5 s for UltrAu grids, blot force of 3 (arbitrary units), wait time of 5 s; chamber temperature of 6 °C, and relative humidity of 95%. Grids were plunge-frozen in liquid ethane cooled by liquid nitrogen.

Cryo-EM data were collected in automated multi-hole mode on a 300 kV Titian Krios microscope for both 1-nt and 10-nt gapped DNA samples, controlled by SerialEM^67^. Micrographs were recorded at a nominal magnification of 105,000× on a K3 direct electron detector (Gatan) operated in super-resolution counting mode, with the objective lens underfocused in the range of 1.2–1.6 μm. For the 1-nt gapped DNA sample, 111-frame movie stacks were recorded with a total exposure time of 3.4 s and a cumulative electron dose of 49.6 e^-^/Å^2^. For the 10-nt gapped DNA sample, 50-frame movie stacks were recorded in a total exposure time of 1 s and a cumulative dose of 59 e^-^/Å^2^. The calibrated physical pixel size at the specimen level was 0.828 Å. Additional data collection parameters are summarized in **Tables S1** and **S2**.

### Cryo-EM data processing

Two cryo-EM datasets comprising 16,140 and 27,515 raw micrographs were collected for the DnaX-complex–β-clamp bound to 1-nt gapped DNA and 10-nt gapped DNA, respectively. Image quality was monitored during data collection using cryoSPARC Live v4.3.1 and v4.5.3^68^ on a local workstation. Image preprocessing—including patch motion correction (bin 2), contrast transfer function (CTF) estimation and correction, blob-based particle picking (70–150 Å diameter), and particle extraction (bin 4)—were performed concurrently. Following two-dimensional (2D) classification, particle displaying clear structural features were selected and used to train a neural network–based particle picking model in Topaz^72^. After further particle picking, 2D classification, and cleaning, selected particles were merged, and duplicate particles with ≥40% overlap were removed such that only one particle was retained within ≤80 Å distance (**Figures S1**, **S5**).

For the dataset containing 1-nt gapped DNA (16,140 micrographs), approximately 4.1 million particles remained after duplicate removal and were subjected to ab initio 3D reconstruction followed by heterogeneous refinement into four 3D classes. Two classes represented junk volumes and were discarded, whereas the remaining two classes corresponded to two major clamp loading states, termed state 1 and state 2, comprising ∼1.8 and ∼1.7 million particles, respectively. State 1 represents an initial DNA recognition state, whereas state 2 corresponds to a post–DNA-entry state in which the 3′-dsDNA has entered the β-clamp and the clamp gate is variably closed, accompanied by sharp bending of the 5′-towards the 3′-dsDNA bound to the outer surface of the β-clamp (**Figure S1**).

After non-uniform refinement, the maps and associated particles were transferred to RELION 5.0.0 for 3D classification without alignment, followed by focused 3D classification on the DNA region using a mask defined by a red dashed oval shape. Selected classes were then transferred back to cryoSPARC v4.5.3 for global and local CTF refinement to correct aberrations introduced by UltrAu grids, followed by reference-based motion correction and additional non-uniform and local refinement.

Three conformers were resolved for state 1: conformer 1 at 3.14 Å resolution with 105,100 particles (∼6% of the population), in which 9 bp of the 20-bp 5′-dsDNA were ordered at the loader shoulder; conformer 2 at 2.71 Å resolution with 228,387 particles, with 5 bp of the 5′-dsDNA ordered; and conformer 3 at 2.73 Å resolution with 232,057 particles, in which the 5′-dsDNA was fully disordered (**Figures 1** and **S1**). Two conformers were obtained for state 2: conformer 1 at 2.72 Å resolution with 283,092 particles, in which the 3′-dsDNA had entered the β-clamp and the 5′-dsDNA bound the basic patch on the fully open clamp; and conformer 2 at 3.12 Å resolution with 203,554 particles, in which the 3′-dsDNA had progressed further into the clamp, the 5′-dsDNA had dissociated, and the β-clamp gate was narrowed by ∼7 Å relative to conformer 1 (**Figures 2** and **S1**). To enhance the map interpretability, EMReady v2.2.1 was applied to the final maps of all five conformers^79,80^.

For the dataset containing 10-nt gapped DNA (27,515 micrographs), approximately 8.5 million particles were used for ab initio 3D reconstruction, followed by heterogeneous refinement into three classes in cryoSPARC. The first class lacked discernible structural feature and was discarded. The second class, containing ∼3.1 million particles, corresponded to the DnaX-complex alone and was subjected to two additional rounds of 2D classification. Selected particles were further refined by heterogeneous refinement using three copies of the initial map as references. One resulting class containing 478,123 particles exhibited the best structural features and was further refined by reference-based motion correction and non-uniform refinement to yield a final map at 2.95 Å resolution (**Figure S5**).

The third initial class, comprising ∼4.7 million particles, corresponded to the ternary DnaX-complex–β-clamp–10-nt gapped DNA complex. This class underwent an additional round of heterogeneous refinement using eight copies of the initial map as references. Of the resulting eight classes, four were discarded due to low resolution (4–7 Å). The remaining four classes, resolved at 3.38–3.57 Å, represented distinct clamp loading intermediates: (1) a DNA gap recognition state in which the 3′-dsDNA had entered the loader chamber but its blunt end remained outside the β-clamp; (2) a state in which the 3′-dsDNA occupied both the loader and clamp chambers with the clamp gate fully open; (3-4) states in which the β-clamp gate was fully closed but exhibited variable flexibility.

Following additional 3D classification, merging of similar classes, reference-based motion correction, and non-uniform refinement, seven conformers of the DnaX-complex–β-clamp–10-nt gapped DNA were obtained: conformer 1 with 473,295 particles at 2.54 Å resolution (DNA recognition state); conformer 2 with 502,506 particles at 2.54 Å resolution (fully open, DNA unsettled); conformer 3 with 499,551 particles at 2.53 Å resolution (fully open, DNA settled); conformer 4 with 476,182 particles at 2.60 Å resolution (partially closed); and conformers 5–7 with 360,948, 539,340, 443,597 particles at 2.88, 2.60 and 2.70 Å resolution, respectively, in which the β-clamp gate was fully closed but the interfaces between DnaX-complex and β-clamp varied (**Figure S5**).

The clamp loader used for cryo-EM analysis was the DnaX-holo-complex containing both ψ and χ subunits. These subunits are small (137 and 147 amino acids; 15.2 and 16.6 kDa, respectively). and diffuse density corresponding to χψ was observed near the expected binding site in some 2D class averages. To improve visualization of this region, all χψ-containing particles (∼7.8 million) from datasets containing the loader alone and the loader bound to β-clamp and 10-nt gapped DNA were combined after heterogeneous refinement and converted to RELION 5.0.0^81^ using PyEM^82^. Focused 3D classification without alignment was performed using a mask surrounding the χψ density. One class containing 242,505 particles at 7.10 Å resolution exhibited clear density for the χψ heterodimer and was transferred back to cryoSPARC for local refinement, yielding a map at 5.50 Å resolution (**Figure S5**). This refined map was merged with the DNA gap recognition state map to generate a composite structure of the χψ-containing DnaX-holo-complex engaged in β-clamp loading onto 10-nt gapped DNA.

### Model building, refinement, and validation

Crystal structures of the γ-complex (PDB 1JR3)^7^, the β-clamp–DNA complex (PDB 3BEP)^83^, and SSB peptide–bound χ–ψ subcomplex (PDB 3SXU)^36^ were docked into the composite cryo-EM map of the DnaX-holo-complex bound to β-clamp and 10-nt gapped DNA at the first loading state (DNA gap recognition). The docked components were merged into a single model using ChimeraX^75^ and served as the initial model for subsequent refinement. The automated *de novo* model-building program ModelAngelo^84^ was also used to facilitate model construction.

Following manual refinement in COOT^76^, the model was subjected to real space refinement in PHENIX^77^, followed by two additional rounds of iterative refinement between PHENIX and COOT. ISOLDE, a molecular-dynamics-based flexible fitting-based plugin within ChimeraX, was then used to minimize model clashes and improve local geometry. The final model was validated using the MolProbity program as implemented in PHENIX^85^.

This refined model was subsequently used as the starting model for building atomic models corresponding to the five loading conformers with 1-nt gapped DNA and the seven loading conformers with 10-nt gapped DNA. In regions of weaker densities, such as near the open gate of the β-clamp, the model was rigid-body docked, followed by manual adjustment in COOT and final real space refinement in PHENIX. All models were refined using the same procedure. Statistics for model refinement and validation are provided in **Tables S1 and S2**. Structural panels were prepared using ChimeraX^86^ and assembled into figures in Adobe Illustrator (Adobe Inc, San Jose, CA).

### Fluorescence polarization–based DNA binding assay

The affinity of the γ-complex for DNA was measured using either a tailed DNA substrate (primer/template, p/T) or gapped DNA substrates containing 10- or 30-nt gaps, at a constant DNA concentration of 10 nM. Importantly, γ-complex was used instead of τ-complex, since τ has a non-specific DNA binding affinity for ssDNA and dsDNA. DNA oligonucleotides were gel purified prior to use. DNA substrates were generated by annealing a 3′ Cy3-labelled Primer 1 either to a 50-nt template (p/T DNA), to Primer 2 plus a 50-nt template (10-nt gapped DNA), or to Primer 2 plus 70-nt template (30-nt gaped DNA) (**Figure 6B**), The annealed substrates were purified on a 10% native polyacrylamide gel. Corresponding gel bands were excised, and DNA was extracted in TE buffer (10 mM Tris-HCl, pH 7.5, 1 mM EDTA), followed by concentration and buffer exchange into TE.

Fluorescence polarization–based binding assays were performed by titrating the γ-complex from 0 to 600 nM into reactions into 50-μL reactions containing 20 mM Tris-HCl (pH 7.5), 40 mM NaCl, 8 mM MgCl_2_, 1 mM TCEP, 0.1 mM EDTA, and 100 μM ATPγS. Reactions were assembled in a 384-well plate and incubated at 23 °C for 20 min. Plates were centrifuged for 20 s to remove air bubbles, and fluorescence signals were recorded using a Synergy Neo2 plate reader (Agilent, Sant Clara, CA), with excitation at 535 nm and collection of emission between 560 and 580 nm. Three independent experiments were performed for each titration.

Relative fluorescence intensity (I/I_0_) was plotted as a function of γ-complex concentration. Apparent dissociation constants (^App^*K_d_*) were determined by fitting the data to a one-site binding model using Origin Software (OriginLab, Northampton, MA).

### Measurement of clamp loading rate onto primed versus gapped DNA

The rate of clamp loading was measured using a replication assay with Pol III αε complex, which requires a loaded β-clamp to extend a primed DNA substrate (**Figures 6C** and **S8A**). In this assay, β-clamp was loaded onto a 5′ ^32^P-labeled primed site in the presence of γ-complex and Pol III αε complex, and timed aliquots were removed for analysis by urea–PAGE. To ensure that clamp loading was the rate-limiting step, Pol III was supplied in sufficient excess such that the distribution of reaction products was determined by clamp loading rather than by the rate of primer extension. Under these conditions, DNA products at all time points exhibit the same size distribution on urea–PAGE but increase in intensity over time. As a control, reactions containing Pol III alone (lacking β-clamp and γ-complex) were performed and showed no primer extension, as expected.

The DNA template (replication template) and the 5′ ^32^P-labeled primer (replication primer 1) used in these assays are listed in the **KEY RESOURCES TABLE**. For gapped DNA substrates, a second oligonucleotide (replication primer 2) was annealed downstream of the 3′ terminus of the 5′ ^32^P-labeled replication primer 1 on a 45-nt template. The replication template strand contained biotin at both termini to retain loaded clamps. For singly primed DNA substrates, replication primer 1 was labeled with ^32^P using T4 polynucleotide kinase (Sigma, St. Louis, MO) and annealed to the replication template. For the 45-nt gapped template, both the ^32^P-labeled replication primer 1 and the unlabeled replication primer 2 were annealed to the replication template.

Reactions were assembled in a scaled-up volume (4.5×; 90 μL total) to allow removal of multiple timed aliquots. Each reaction contained 0.9 pmol of ^32^P-primed template and 5.3 pmol streptavidin tetramer in Buffer B supplemented with 8 mM MgCl_2_. Reactions were incubated at 23 °C for 15 min, followed by addition of 2.7 pmol SSB and incubation for an additional 5 min at 37 °C. Reactions were initiated by addition of 0.9 pmol *E. coli* Pol III, 0.9 pmol β-clamp, and 0.9 pmol γ-complex in a final volume of 90 μL buffer B containing 8 mM MgCl_2_, 2 mM ATP, and 100 mM each of dATP, dCTP, dGTP and ^32^P-dTTP.

Reactions were incubated at 23 °C, and 20-μL aliquots were withdrawn at 0.5, 1, 2, and 5 min and quenched with 25 μL stop solution (76% formamide, 14 mM EDTA, 0.02% bromophenol blue, 2% SDS). Replicated products were resolved on a 15% urea–denaturing polyacrylamide gel (**Figure S8A**). Gels were exposed to a storage phosphor screen and scanned using a Typhoon PhosphorImager (GE Healthcare). Product bands were quantified using ImageQuant software (GE Healthcare). Each reaction was performed in triplicate, and mean values and standard deviations were calculated using Microsoft Excel.

### ATPase assays

Two oligonucleotides (oligo 1: 5’-GGCTCGCTACGGCTATACGG-3’; oligo 2: 5’-GATTCGTATCGCCTATACCG-3’) were both annealed to a template oligo, 5’-CGGTATAGGCGATACGAATC(T_n_) CCGTATAGCCGTAGCGAGCC-3’, where n = 0, 1, 2, 5, 10, to generate DNA substrates containing a nick or 1-, 2-, 5- or 10-nt gaps. The resulting DNA substrates were separated from unannealed oligonucleotides by native 20% agarose gel electrophoresis and purified from the gel.

ATPase reactions were performed by mixing 50 nM wild-type γ-complex with 500 nM purified gapped DNA substrates and 200 nM β-clamp in a total volume of 30 μL containing 20 mM Tris-HCl (pH7.5), 0.1 mM EDTA, 40 μg/mL BSA, 4% glycerol, 5 mM DTT, 8 mM MgCl_2_, 0.5 mM ATP, and 55 nM γ-^32^P-ATP. Reactions were incubated at 37 °C, and 5-μL aliquots were removed at 5, 10, 20, and 30 min and quenched with an equal volume of 0.5 M EDTA.

One microliter of each quenched reaction was spotted onto PEI Cellulose F thin-layer chromatography (TLC) plate (Millipore Sigma) and developed in 0.75 M KHPO_4_ (pH 4.4). TLC plates were exposed to a PhosphorImager screen and scanned using an Amersham Typhoon instrument. Radioactive ATP and inorganic phosphate were quantitated using ImageQuant software (**Figure S8B**).

## QUANTIFICATION AND STATISTICAL ANALYSIS

Fluorescence polarization–based binding assays measuring γ-complex interactions with different DNA substrates were performed in three independently performed experiments. Data fitting was carried out using Origin Software (OriginLab, Northampton, MA), and results are presented as mean ± standard error of the (SEM) in **Figure 6B**. Statistical analysis of cryo-EM structures was performed during model refinement in Phenix, with details provided in **Tables S1** and **S2**.

## Supplemental video legends

**Video 1. Overview of thirteen β-clamp loading intermediates captured for DnaX-complex loading β-clamp onto either a 1-nt gapped DNA (five intermediates) or a 10-nt gapped DNA (eight intermediates).** The thirteen atomic models are aligned and displayed individually, each rotated by 360° about the Y axis.

**Video 2. Morphing of β-clamp loading onto a 1-nt gapped DNA substrate.** The morph integrates previously reported structures of DnaX-complex loading β-clamp onto tailed DNA^16,17^, with the five loading intermediates captured here using a 1-nt gapped DNA. The animation begins with formation of a binary DnaX-complex–β-clamp complex in the presence of ATP, which induces β-clamp gate opening. The open clamp then engages the 3’-dsDNA segment of the 1-nt gapped DNA via a conserved basic patch on the outer surface of the β-clamp near the open gate. This interaction is followed by sharp bending of the 5′-dsDNA towards the 3′-dsDNA, generating a stressed U-shaped DNA conformation and exposing the recessed 3′ end. The resulting DNA strain promotes insertion of the recessed 3′ end into the DnaX-complex chamber, entry of the 3′-dsDNA into the β-clamp, and binding of the 5′-dsDNA to the clamp basic patch vacated by the 3’-dsDNA. During this process, the δ-subunit Y316 sensor recognizes—but does not unwind—the recessed DNA 3′ end. Subsequent DNA engagement within the β-clamp chamber is proposed to drive clamp gate closure via electrostatic interactions, stimulate ATP hydrolysis, and trigger dissociation of the DnaX-complex, completing the clamp loading reaction.

**Video 3. Morphing of β-clamp loading onto a larger gapped DNA (>6 nt).** The animation was generated in ChimeraX based on the reported structures of the β-clamp (PDB: 1MMI)^87^, the *E. coli* DnaX-complex bound to β-clamp in the absence of DNA (PDB: 8GIY and 7TI8)^16,17^, and the eight loading intermediates captured here with a 10-nt gapped DNA. In the presence of ATP, the DnaX-complex binds and opens the β-clamp gate. Because the recessed DNA 3′ end is accessible in large-gapped substrates, it can directly enter the clamp loader chamber and engage the δ-subunit Y316 sensor residue, while the 3′-dsDNA binds to the basic patch on the exterior surface of β-clamp. Electrostatic attraction between the negatively charged DNA backbone and the positively charged interior of the β-clamp promotes insertion of the 3′-dsDNA into the clamp chamber and drives clamp gate closure, ultimately leading to dissociation of the DnaX-complex and formation of a DNA-encircling β-clamp.

---

**Table S1.** Cryo-EM data collection, refinement, and atomic model validation.

| Structures | DnaX-complex-β-clamp-1-nt gapped DNA |  |  |  |  | DnaX-complex<sup>a</sup> | ψ-χ focus refined<sup>a</sup> |
| --- | --- | --- | --- | --- | --- | --- | --- |
|  | State 1 conf. 1 | State 1 conf. 2 | State 1 conf. 3 | State 2 conf. 1 | State 2 conf. 2 |  |  |
| EMDB ID | EMD-71013 | EMD-71014 | EMD-71015 | EMD-71016 | EMD-71017 | EMD-71018 | EMD-71020 |
| PDB ID | 9OYB | 9OYC | 9OYD | 9OYE | 9OYF | 9OYG |  |
| **Data collection** |  |  |  |  |  |  |  |
| Magnification | 105,000 |  |  |  |  |  |  |
| Voltage (kV) | 300 |  |  |  |  |  |  |
| Dose (e<sup>-</sup>/Å<sup>2</sup>) | 49.6 |  |  |  |  | 59 |  |
| Under-focus (μm) | 1.2 - 1.6 |  |  |  |  |  |  |
| Pixel size (Å) | 0.828 |  |  |  |  |  |  |
| Symmetry | C1 |  |  |  |  |  |  |
| Initial particle # | 4.1 million |  |  |  |  | 3.1 M | 7.8 M |
| Final particle # | 105,100 | 228,387 | 232,057 | 283,092 | 203,554 | 478,123 | 242,505 |
| Map resolution (Å) | 3.14 | 2.71 | 2.73 | 2.72 | 3.12 | 2.95 | 5.50 |
| FSC threshold |  |  |  |  |  | 0.143 |  |
| Map resolution range (Å) | 2.7 – 11.0 | 2.4 – 11.0 | 2.3 – 12.0 | 2.4 – 10.3 | 2.7 – 10.1 | 2.5 – 9.5 | 4.5 – 13.0 |
| **Refinement** |  |  |  |  |  |  |  |
| Initial model used (PDB code) | 1JR3, 3BEP |  |  |  |  | 1JR3 | 3SXU |
| Map sharpening B factor (Å<sup>2</sup>) | -105.2 | -86.9 | -87.1 | -99.5 | -107.2 | -116.7 | -211.4 |
| Map to model CC<sub>mask</sub> | 0.89 | 0.89 | 0.88 | 0.88 | 0.88 | 0.79 |  |
| **Model composition** |  |  |  |  |  |  |  |
| Non-hydrogen atoms | 21,074 | 20,894 | 20,661 | 21,517 | 21,127 | 14,125 |  |
| Protein and DNA residues | 2,532; 59 | 2,530; 51 | 2,529; 40 | 2,531; 81 | 2,531; 62 | 1794; 0 |  |
| Ligands | 10 | 10 | 10 | 10 | 10 | 10 |  |
| **R.m.s. deviations** |  |  |  |  |  |  |  |
| Bond lengths (Å) | 0.004 | 0.004 | 0.003 | 0.006 | 0.005 | 0.003 |  |
| Bond angles (°) | 0.756 | 0.631 | 0.594 | 0.684 | 0.669 | 0.623 |  |
| **Validation** |  |  |  |  |  |  |  |
| MolProbity score | 1.34 | 1.67 | 1.26 | 1.38 | 1.49 | 1.58 |  |
| Clashscore | 5.53 | 4.56 | 4.99 | 6.08 | 5.47 | 8.42 |  |
| Poor rotamers (%) | 0 | 0 | 0 | 0 | 0 | 0 |  |
| **Ramachandran plot** |  |  |  |  |  |  |  |
| Favored (%) | 97.81 | 98.33 | 98.29 | 97.77 | 96.86 | 97.35 |  |
| Allowed (%) | 2.19 | 1.67 | 1.71 | 2.23 | 3.14 | 2.65 |  |
| Disallowed (%) | 0 | 0 | 0 | 0 | 0 | 0 |  |

<sup>a</sup>: The last two columns are from dataset with 10-nt gapped DNA (continued in **Table S2**).

**Table S2.** Cryo-EM data collection, refinement, and atomic model validation.

| Structures | DnaX-complex-β-clamp-10-nt gapped DNA | | | | | | |
| --- | --- | --- | --- | --- | --- | --- | --- |
|  | DNA recognition | fully open, DNA unsettled | fully open, DNA settled | partially closed | fully closed 1 | fully closed 2 | fully closed 3 |
| EMDB ID | EMD-71021 | EMD-71022 | EMD-71023 | EMD-71024 | EMD-71025 | EMD-71026 | EMD-71027 |
| PDB ID | 9OYH | 9OYI | 9OYJ | 9OYK | 9OYL | 9OYM | 9OYN |
| **Data collection and processing** |  |  |  |  |  |  |  |
| Magnification | 105,000 |  |  |  |  |  |  |
| Voltage (kV) | 300 |  |  |  |  |  |  |
| Electron dose (e<sup>-</sup>/Å<sup>2</sup>) | 59 |  |  |  |  |  |  |
| Under-focus range (μm) | 1.2 - 1.6 |  |  |  |  |  |  |
| Pixel size (Å) | 0.828 |  |  |  |  |  |  |
| Symmetry imposed | C1 |  |  |  |  |  |  |
| Initial particle images (no.) | 8,537,193 |  |  |  |  |  |  |
| Final particle images (no.) | 473,295 | 502,506 | 499,551 | 476,182 | 360,948 | 539,340 | 443,597 |
| Map resolution (Å) | 2.54 | 2.54 | 2.53 | 2.60 | 2.88 | 2.60 | 2.70 |
| FSC threshold | 0.143 |  |  |  |  |  |  |
| Map resolution range (Å) | 2.2 – 9.5 | 2.2 – 9.3 | 2.2 – 9.5 | 2.3 – 9.6 | 1.8 – 9.9 | 2.3 – 9.4 | 2.4 – 9.9 |
| **Refinement** |  |  |  |  |  |  |  |
| Initial model used (PDB code) | 1JR3, 3BEP, 3SXU |  |  |  |  |  |  |
| Map sharpening B factor (Å<sup>2</sup>) | -91.6 | -92.5 | -92.6 | -95.7 | -99.1 | -98.1 | -99.0 |
| Map to model CC<sub>mask</sub> | 0.71 | 0.87 | 0.75 | 0.87 | 0.75 | 0.86 | 0.82 |
| **Model composition** |  |  |  |  |  |  |  |
| Non-hydrogen atoms | 22,702 | 20,796 | 20,835 | 20,861 | 20,835 | 20,861 | 20,843 |
| Protein and DNA residues | 2,779; 44 | 2,531; 46 | 2,531; 47 | 2,534; 47 | 2,532; 47 | 2,534; 47 | 2,532; 47 |
| Ligands | 10 | 10 | 11 | 11 | 11 | 11 | 11 |
| **R.m.s. deviations** |  |  |  |  |  |  |  |
| Bond lengths (Å) | 0.002 | 0.005 | 0.007 | 0.003 | 0.003 | 0.004 | 0.005 |
| Bond angles (°) | 0.480 | 0.634 | 0.687 | 0.530 | 0.489 | 0.587 | 0.579 |
| **Validation** |  |  |  |  |  |  |  |
| MolProbity score | 1.40 | 1.21 | 1.52 | 1.24 | 1.45 | 1.21 | 1.37 |
| Clashscore | 7.29 | 4.31 | 6.67 | 4.73 | 6.98 | 4.25 | 5.19 |
| Poor rotamers (%) | 0 | 0 | 0 | 0 | 0 | 0 | 0 |
| Ramachandran plot |  |  |  |  |  |  |  |
| Favored (%) | 98.15 | 98.17 | 97.13 | 98.13 | 97.69 | 98.57 | 97.54 |
| Allowed (%) | 1.85 | 1.83 | 2.87 | 1.87 | 2.31 | 1.43 | 2.46 |
| Disallowed (%) | 0 | 0 | 0 | 0 | 0 | 0 | 0 |

**Figure S1.**
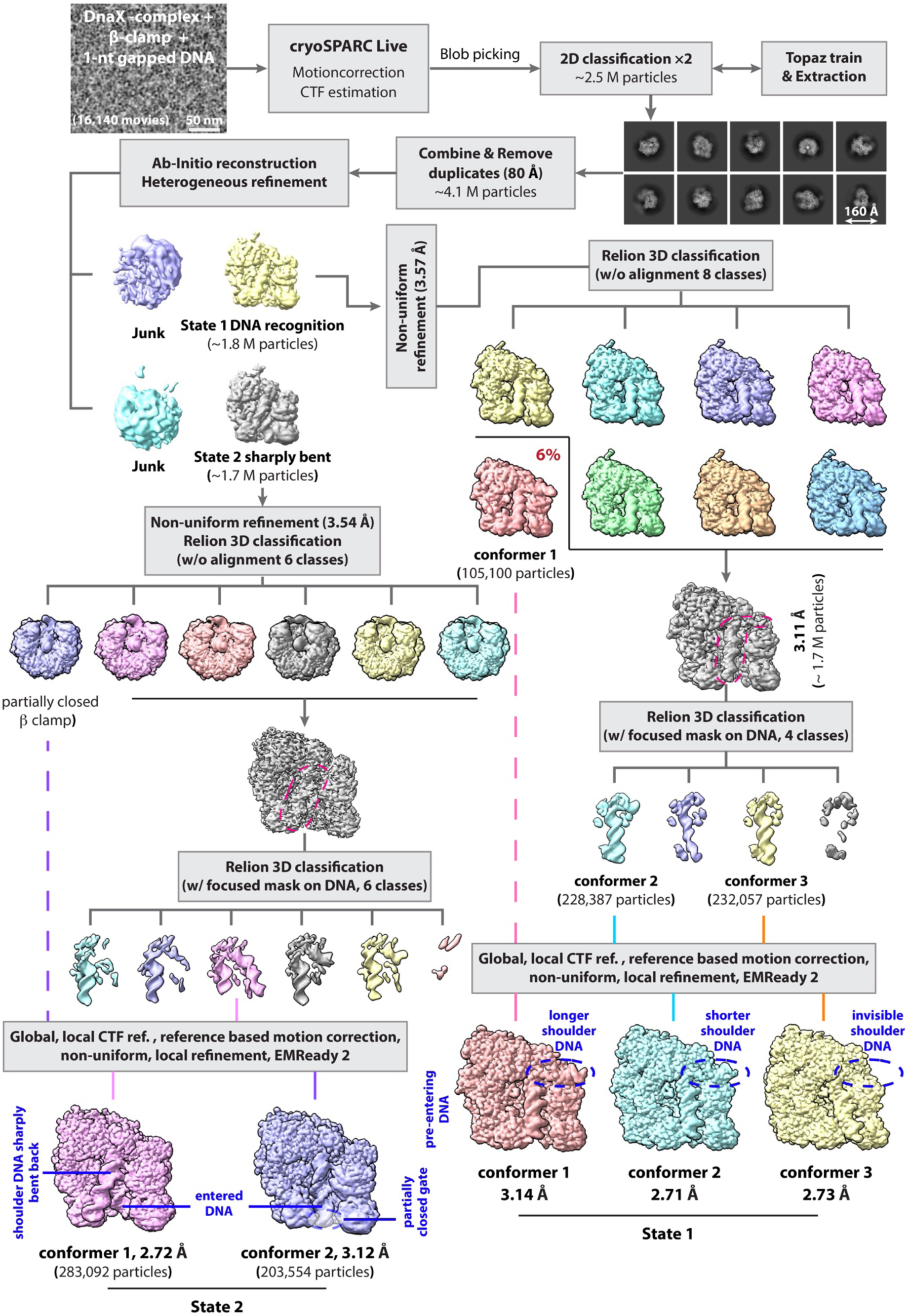
Cryo-EM data processing of the DnaX-complex−β-clamp bound to 1-nt gapped DNA. A representative micrograph from approximately 16,000 movies collected on a 300 kV Titan Krios microscope is shown, with scale bar indicated. Image processing was performed using cryoSPARC v4.5 and Relion v5.0^68,88^. Two major clamp loading states were identified: a first state comprising three conformers and a second state comprising two conformers.

**Figure S2.**
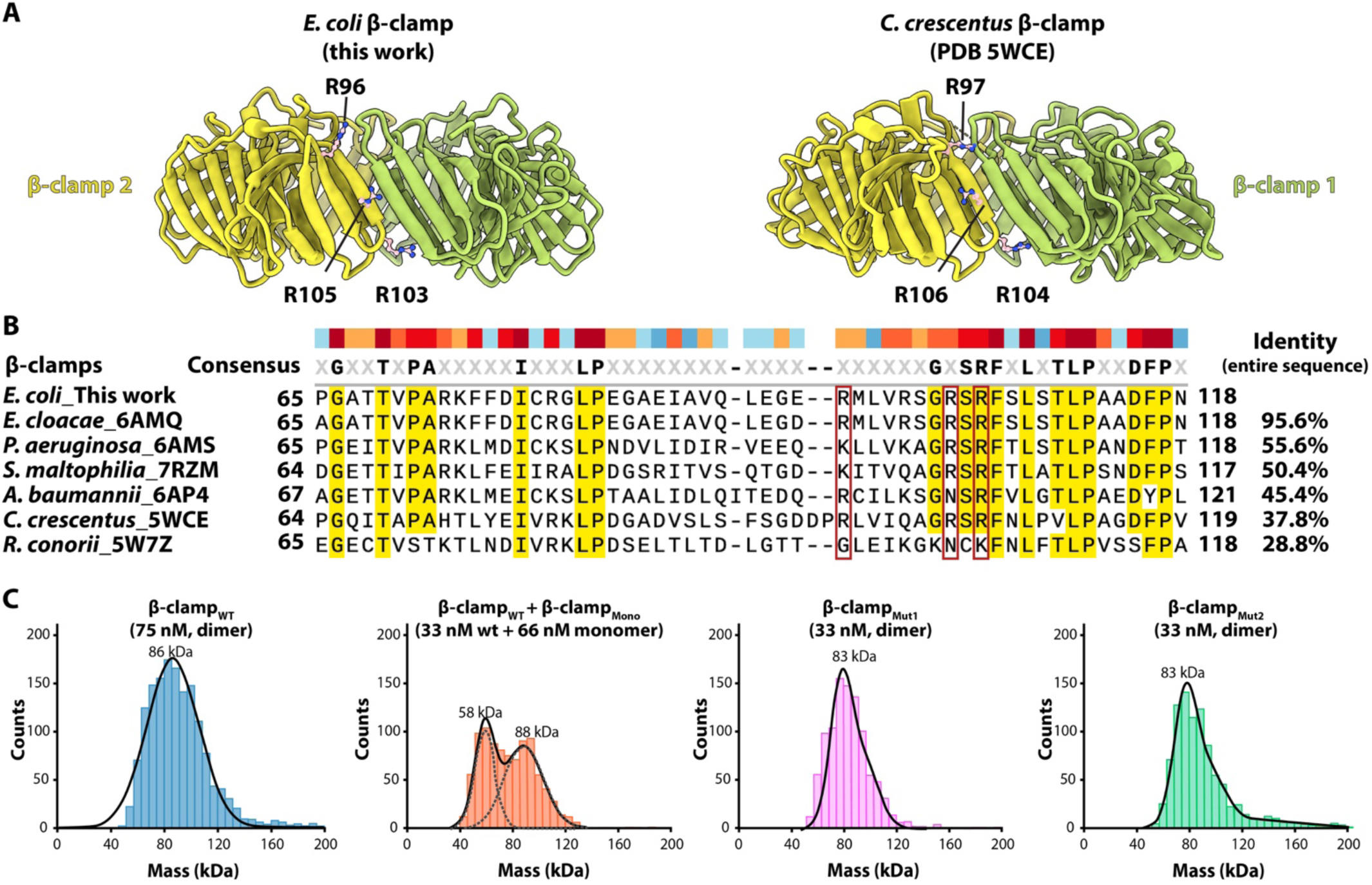
The basic patch on the outer surface of the β-clamp is conserved among prokaryotes. Related to Figure 3. (**A**) Structural comparison of the *E. coli* β-clamp with that of *Caulobacter crescentus* as a representative example. The three positively charged residues that form the conserved basic patch are shown as sticks and are located on the outer surface near the left side of the DNA entry gate between the two β subunits. (**B**) Structure-based sequence alignment of the β-clamp region containing the three basic residues, highlighted by red boxes. Structures were aligned using ChimeraX^86^, and PDB accession code for each structure is listed following the species name. (**C**) The β-clamp mutants used in this study remain dimeric in solution. Mass photometry measurements have a low molecular weight detection limit of ∼40 kDa; therefore, apparent β monomer mass values are expected to be slightly higher than theoretical monomer mass. As controls, wild-type β-clamp dimers and a previously characterized β monomer mutant, in which the hydrophobic core residues were mutated to disrupt dimerization (I272A/L273/A)^89^ were analyzed. Both β-clamp_Mut1_ and β-clamp_Mut2_, harboring R96, R103, and R105 substitutions to alanine or glutamate, respectively, remained dimeric in solution.

**Figure S3.**
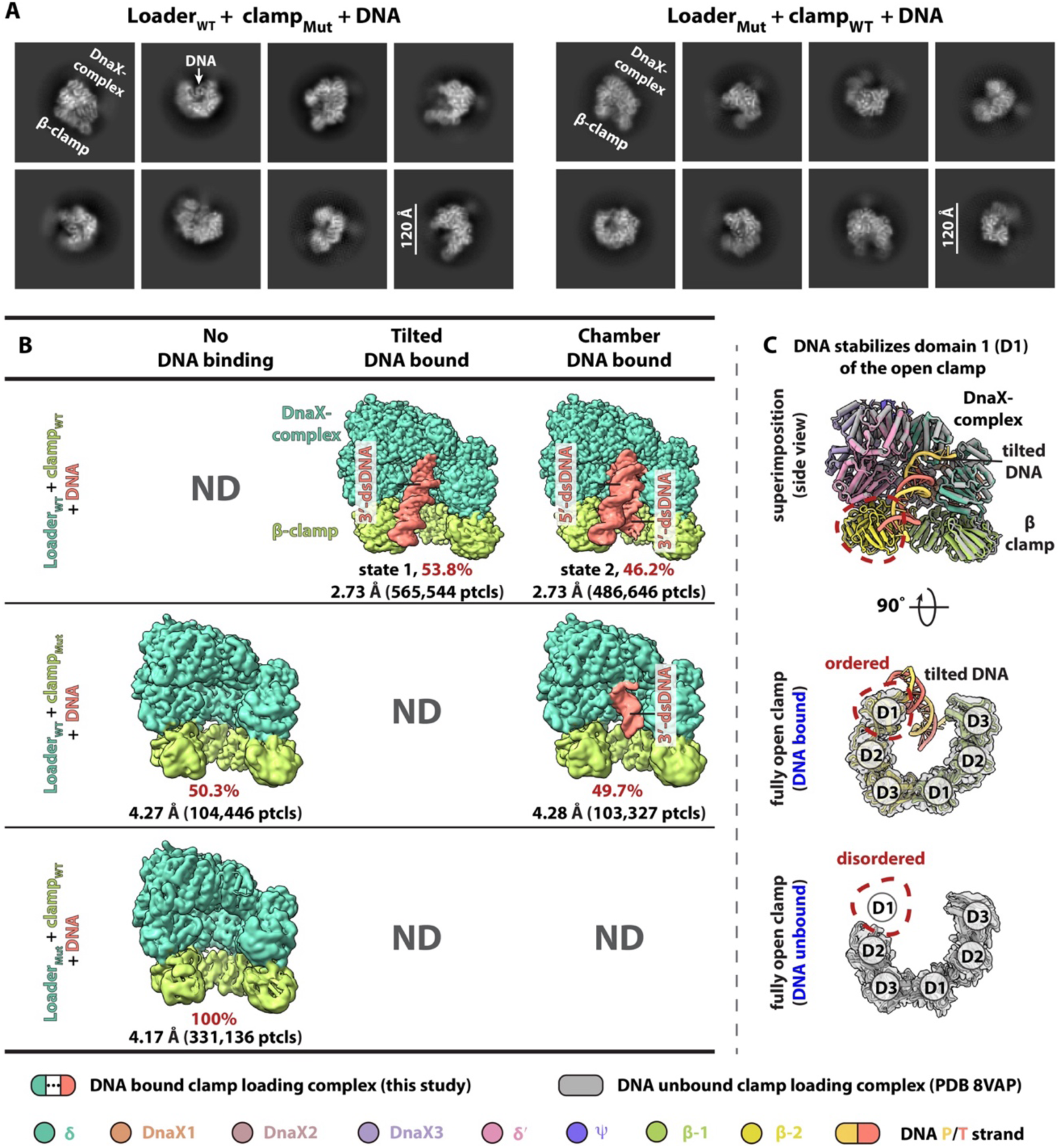
Cryo-EM analysis of the effects of mutations on β-clamp loading. (**A**) Representative 2D class averages of assembled clamp loading complexes containing mutant components. Left: wild-type DnaX-complex (loader_WT_) assembled with a β-clamp basic patch mutant harboring R96A, R103A, and R105A substitutions (β-clamp_Mut_) and 1-nt gapped DNA. Right: DnaX-complex bearing the δ-subunit Y316A substitution (loader_Mut_) assembled with wild-type β-clamp (clamp_WT_) and 1-nt gapped DNA. Assemblies were performed under same conditions as used for wild-type complexes. Key components and scale bar are indicated. (**B**) Reconstructed cryo-EM maps corresponding to the assemblies shown in (A). For comparison, representative maps from wild-type clamp and clamp loader assemblies are also shown (state 1 conformer 1 and state 2 conformer 1). Total particle numbers for each state (summed across conformers) are listed to indicate relative populations. For assemblies containing wild-type clamp and loader, no clamp–clamp loader–only complex (lacking DNA) was observed. In contrast, assemblies containing β-clamp_Mut_ yielded approximately equal populations of clamp–clamp loader–only complexes and complexes containing chamber-bound 3′-dsDNA, but no intermediates with tilted 3′-dsDNA or bent 5′-were detected, consistent with the requirement for the β-clamp basic patch in DNA binding. For assemblies containing loader_Mut_, only clamp–clamp loader–only complex was observed, with no DNA - bound state observed, indicating that the δ-subunit Y316 is critical for DNA recognition and subsequent clamp loading. ND, not detected. (**C**) DNA binding at the β-clamp basic patch stabilizes β-clamp domain I. Superimposition of structures of the DnaX-complex with an open β-clamp in the absence of DNA (gray, PDB: 8VAP)^17^ and in the presence of 1-nt gapped DNA (colored; state 1 conformer 1 from this study) is showed. The clamp loader is omitted in the bottom views for clarity.

**Figure S4.**
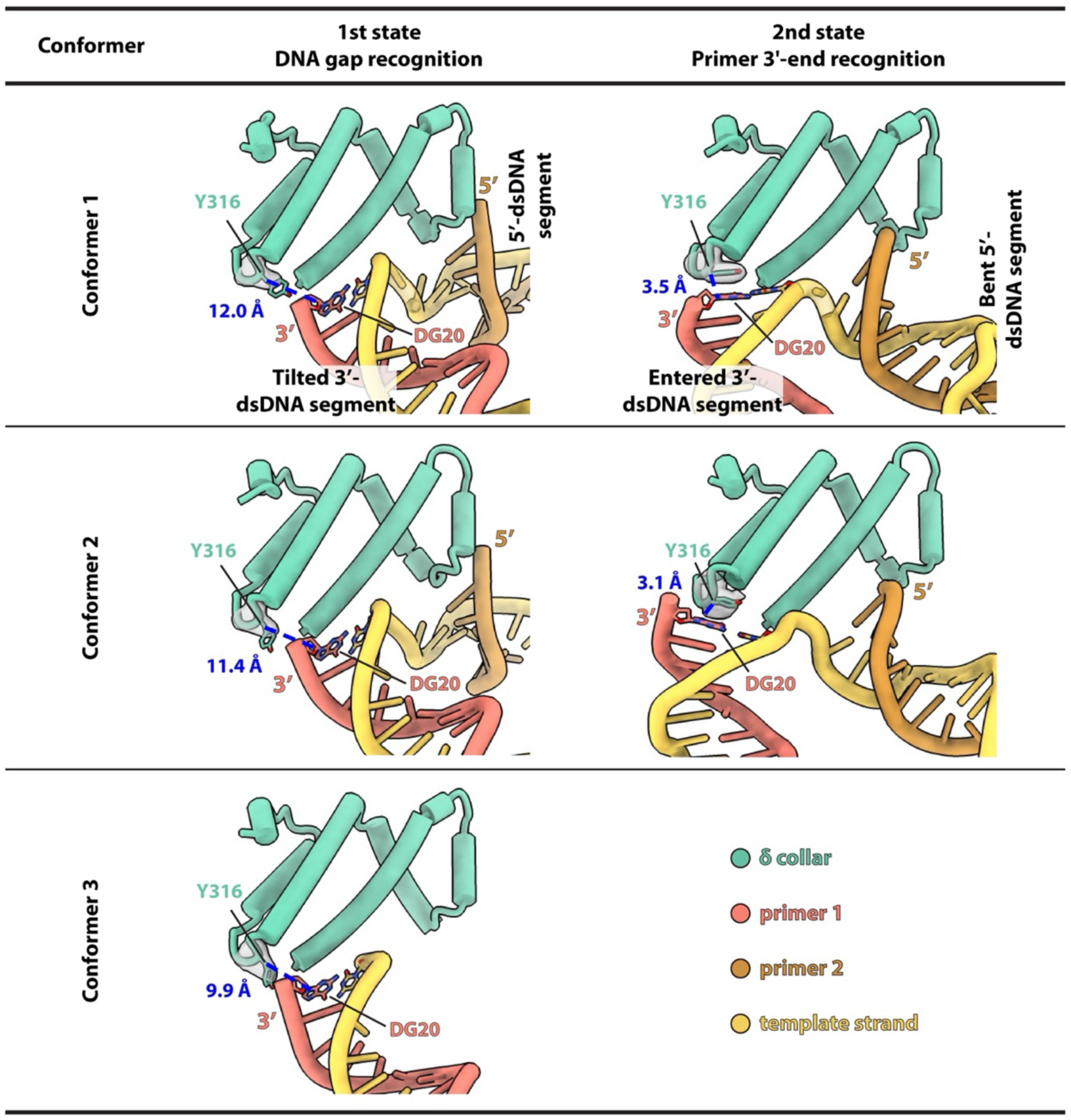
The *E. coli* clamp loader residue δ Y316 recognizes the recessed DNA 3′ end to facilitate proper positioning of the 3′-dsDNA within the loader chamber. In the first DNA gap recognition state, comparison of conformers 1–3 shows a progressive decrease in the distance between δ Y316 and the DNA 3′ end, from 12.0 Å to 9.9 Å, accompanied by increasingly well-defined local cryo-EM density for Y316. In the second state, corresponding to DNA primer 3′ end recognition, this distance is further reduced to ∼3.3 Å in both conformers, and Y316 adopts a base-mimicking conformation that stacks against the terminal base G20 of the primer 3′ end, resulting in full stabilization of this residue. For clarity, only a closed-up view of this region is shown. Distance were measured between the Y316 Cβ atom and G20 C8 atom, which approximates the distance between the centers of the two ring-containing residues in the second state.

**Figure S5.**
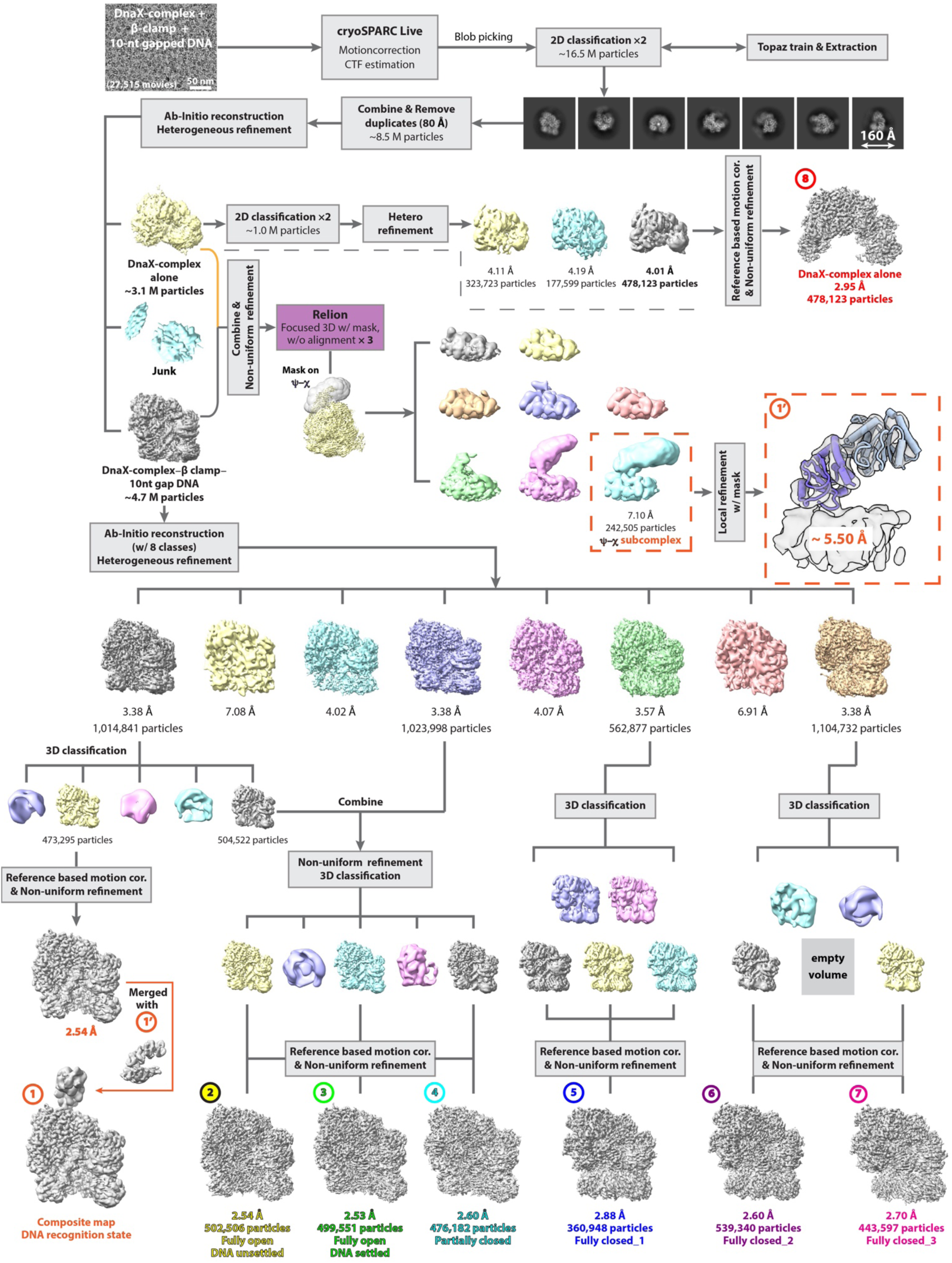
Cryo-EM data processing of the DnaX-complex−β-clamp bound to 10-nt gapped DNA. A representative micrograph from ∼28,000 movies collected on a 300kV Titan Krios microscope is shown, with scale bar indicated. Image processing was performed using cryoSPARC v4.5 and Relion v5.0^68,88^. Eight distinct clamp loading states were resolved at resolutions from ∼3.0 Å to 2.5 Å, with density corresponding to the χψ heterodimer merged into the state 2 map. The χψ crystal structure bound to an SSB peptide bound (PDB: 3SXU) was rigid-body fitted into the locally refined 3D map highlighted by dashed red squares.

**Figure S6.**
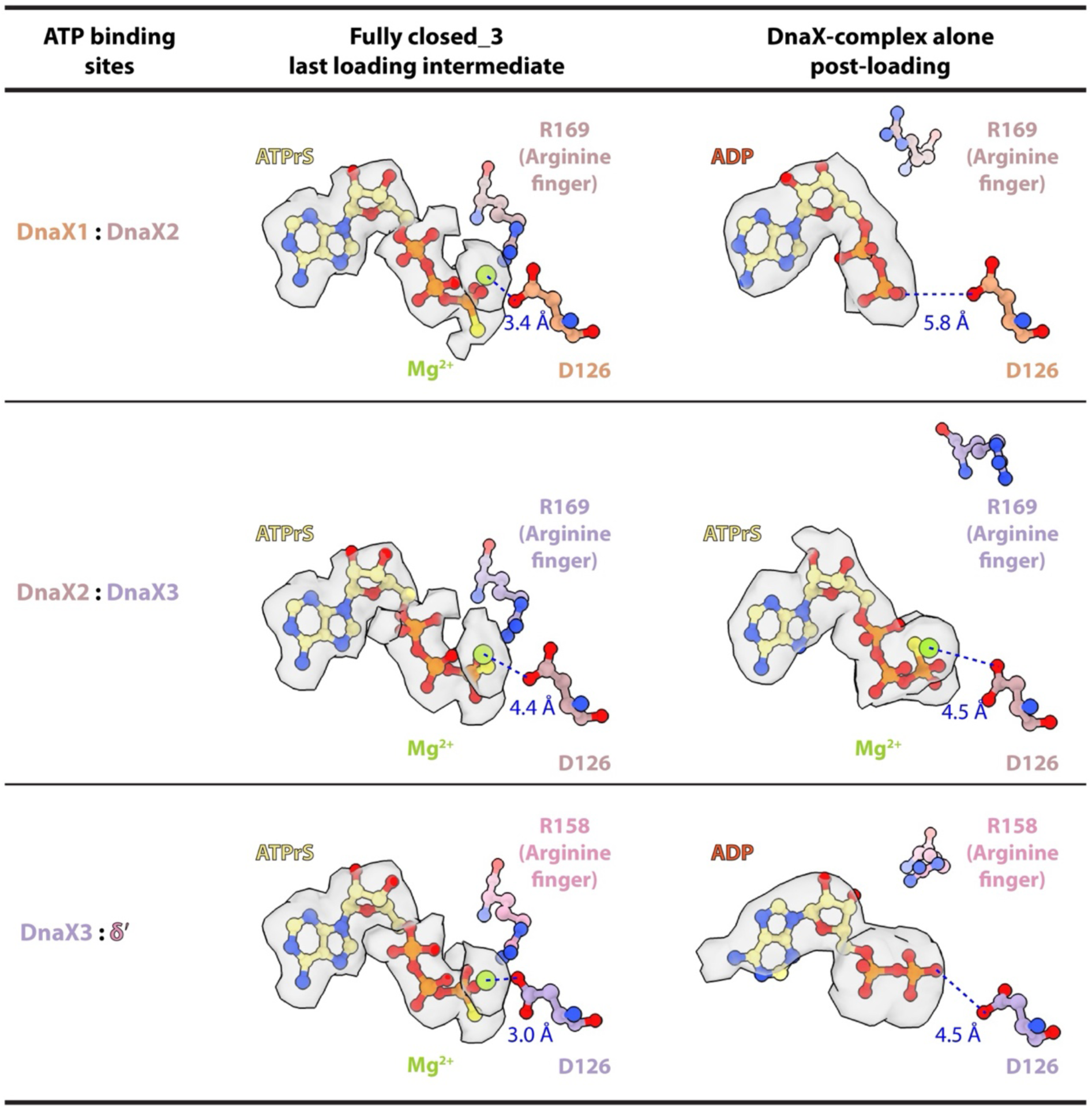
ATPγS hydrolysis is observed only in the DnaX-complex–alone structure. Across all 13 clamp loading intermediates captured with either 1-nt or 10-nt gapped DNA, ATPγS was bound to all ATPase sites from the fully open β-clamp state through the final fully closed state, indicating no ATPγS hydrolysis during these intermediates. The last loading intermediates (fully closed_3) is shown here as a representative example. In this state, the distances between the Mg^2+^ ion and the catalytic residue D126 (Walker B/DEAD box) are 3.4 Å and 3.0 Å at the first and third ATPase sites, respectively, consistent with a geometry competent for hydrolysis, whereas the distance at the second site is 4.4 Å. By contrast, in the DnaX-complex–alone structure, ADP can be unambiguously modeled at the first and third sites, while ATPγS remained bound at the second site, indicating that ATPγS hydrolysis had occurred at two sites following clamp loading. For clarity, only the bound nucleotides and key catalytic residues are shown.

**Figure S7.**
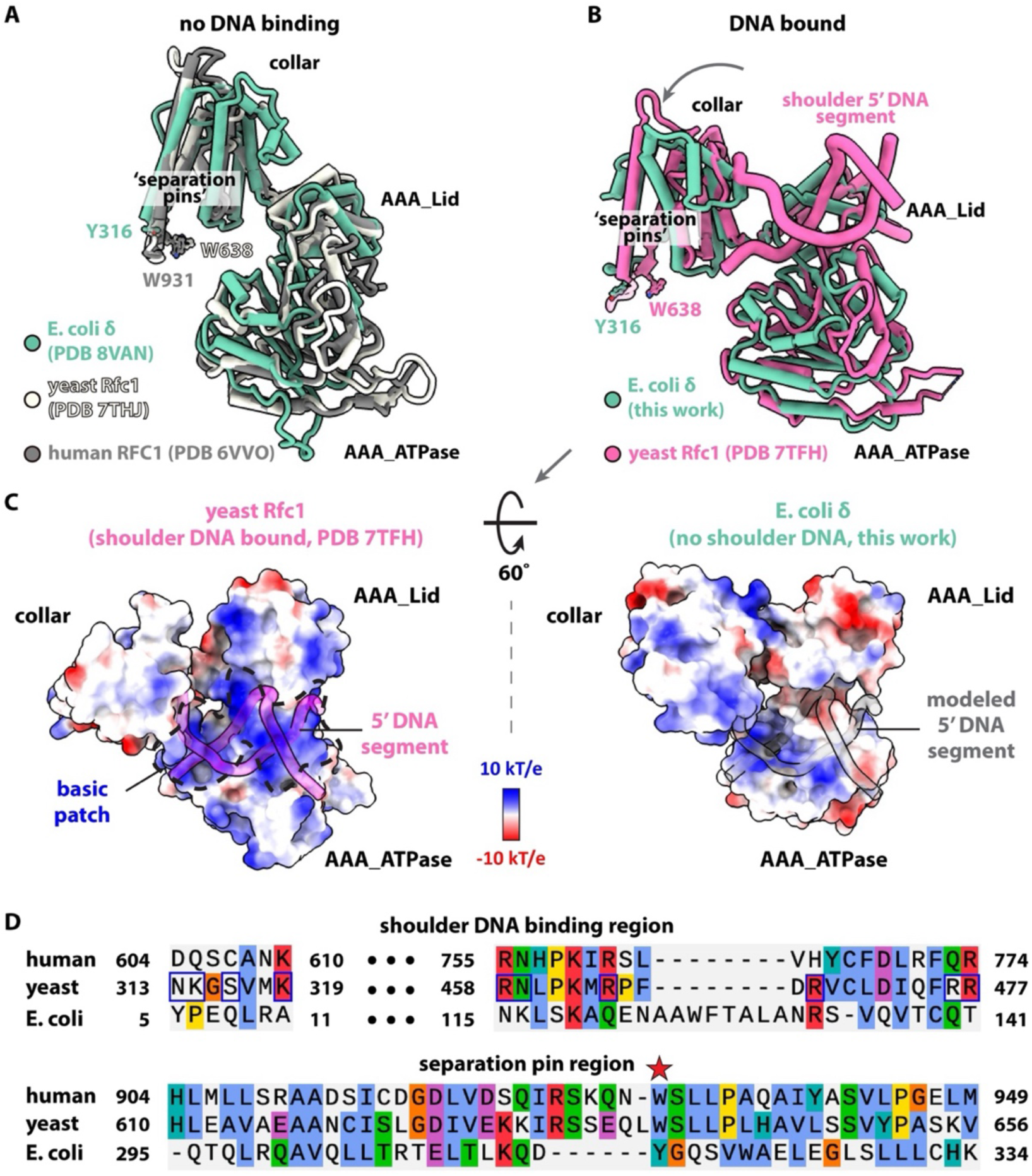
Comparisons of the clamp loader first subunit from *E. coli*, yeast, and human. (**A**) In the absence of DNA binding, the overall structures of *E. coli* δ, yeast Rfc1, and human RFC1 are similar. (**B**) Upon binding of the 3′-dsDNA in the chamber (omitted), the yeast Rfc1 AAA Lid and Collar domains undergo a larger movement than those in *E. coli*, creating additional space to facilitate subsequent binding of 5′-dsDNA at the shoulder. (**C**) The Rfc1 shoulder region presents a large basic patch (black dashed line) for 5′-dsDNA binding (left), which is absent in *E. coli* δ; steric clashes are observed when the Rfc1-bound 5′-dsDNA is modeled onto *E. coli* δ (right). (**D**) Structural-based sequence alignment shows that the shoulder regions of yeast Rfc1 and human RFC1 are more positively charged than that of *E. coli* δ (upper). The ‘separation pin’ in human and yeast proteins is a Trp residue with two aromatic rings, whereas the *E. coli* protein contains a Tyr residue with a single aromatic ring (lower). Residues are colored by chemical property (scheme from the MUSCLE alignment program^90^).

**Figure S8.**
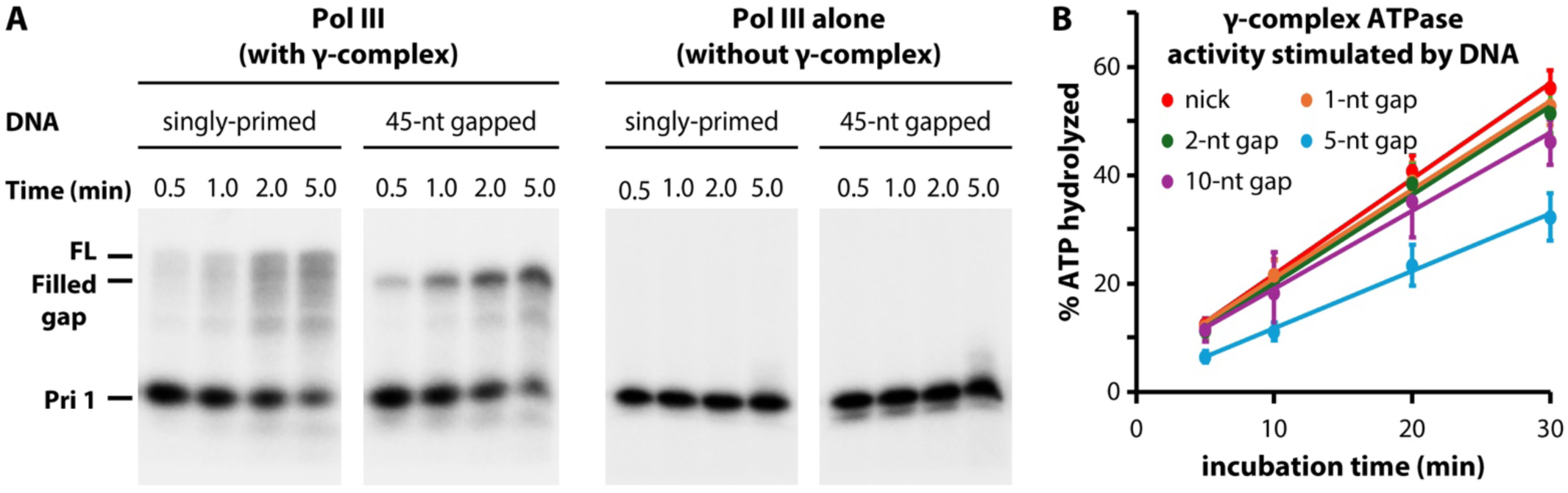
Characterization of Pol III DNA synthesis and DnaX-complex ATPase activity on distinct DNA substrates. (**A**) Urea–PAGE analysis of ^32^P-labelled primer extension over increasing loading times using either singly primed or gapped DNA, in the presence (left) or absence (right) of the γ-complex. A sufficient amount of Pol III was added to saturate the reaction, making clamp loading the rate limiting step. (**B**) Stimulation of DnaX-complex ATPase activities by DNA substrates containing gaps of different sizes. See **Methods** for details.

**Figure S9.**
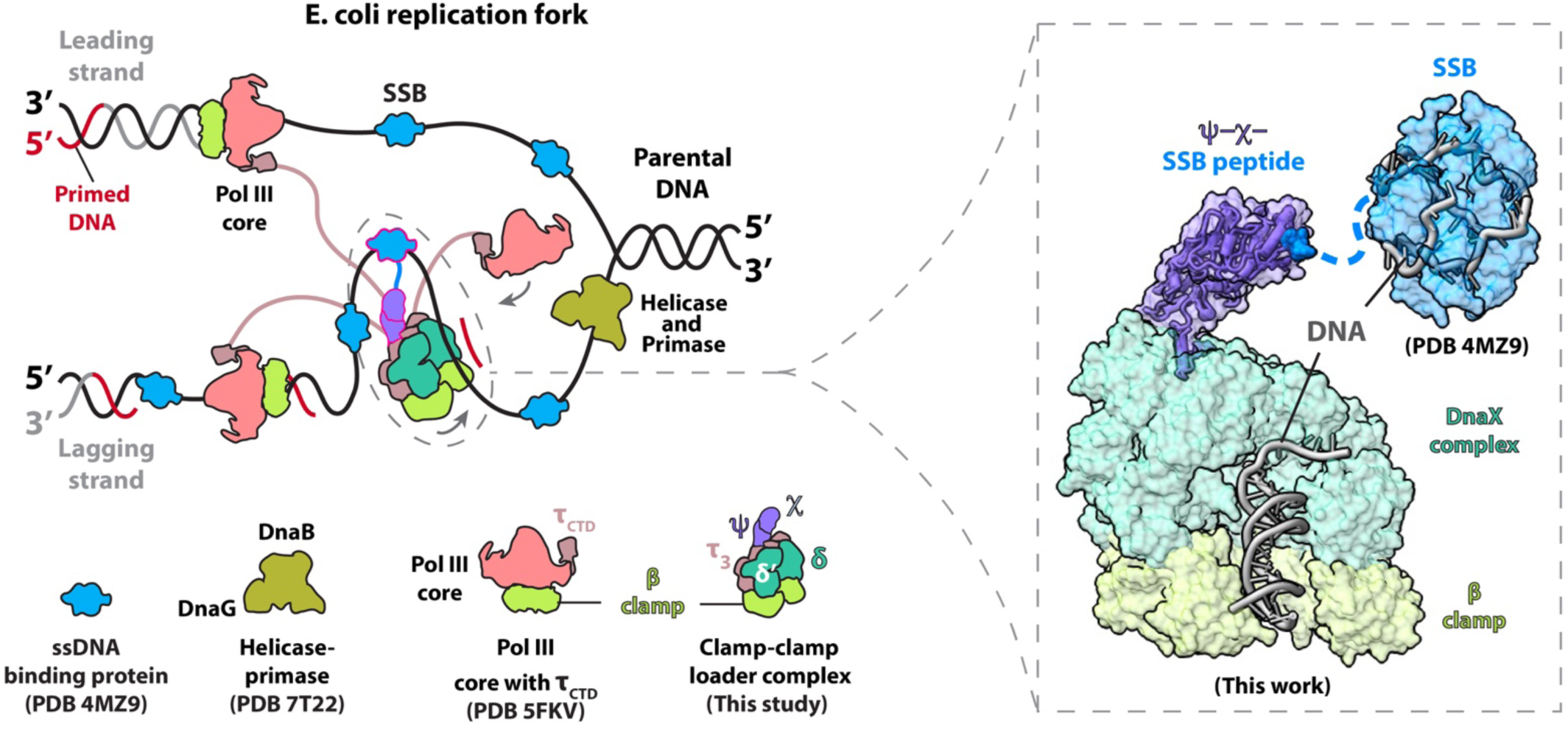
Model for coordination of the replisome by the DnaX-holo-complex. Left, a simplified model of the *E. coli* replication fork is shown, including the DnaB helicase encircling the lagging strand and unwinding parental DNA, the DnaG primase bound to DnaB and synthesizing RNA primers on the lagging strand template, and multiple SSB molecules coating the ssDNA. In addition to loading the β-clamp for Pol III οn both the leading and lagging strands, the DnaX-holo-complex organizes the replication fork by interacting with SSB via its χ subunit and with three Pol III molecules via the C-terminal domains of its three τ subunits (τ_CTD_)^61,62^. These interactions may facilitate recruitment of Pol III to the loaded β-clamp. Key components are listed together with the available PDB codes. Right, a structural model shows SSB bound to ssDNA and interacting with the DnaX-holo-complex through its flexibly linked C-terminal peptide (blue surface).

## Notes

### Competing Interest Statement

The authors have declared no competing interest.

### Summary of Updates

Added more biochemical and structural characterizations of the clamp loading system with various gap sizes (Figure 3A, Figure S3, S4, S5, S8).

## REFERENCES

1. Stukenberg, P.T., Studwell-Vaughan, P.S., and O’Donnell, M. (1991). Mechanism of the sliding beta-clamp of DNA polymerase III holoenzyme. The Journal of biological chemistry 266, 11328–11334.

2. Kong, X.P., Onrust, R., O’Donnell, M., and Kuriyan, J. (1992). Three-dimensional structure of the beta subunit of E. coli DNA polymerase III holoenzyme: a sliding DNA clamp. Cell 69, 425–437. 10.1016/0092-8674(92)90445-i.

3. Kelch, B.A., Makino, D.L., O’Donnell, M., and Kuriyan, J. (2012). Clamp loader ATPases and the evolution of DNA replication machinery. BMC Biol 10, 34. 10.1186/1741-7007-10-34.

4. O’Donnell, M., Langston, L., and Stillman, B. (2013). Principles and concepts of DNA replication in bacteria, archaea, and eukarya. Cold Spring Harb Perspect Biol 5. 10.1101/cshperspect.a010108.

5. Krishna, T.S., Kong, X.P., Gary, S., Burgers, P.M., and Kuriyan, J. (1994). Crystal structure of the eukaryotic DNA polymerase processivity factor PCNA. Cell 79, 1233–1243. 10.1016/0092-8674(94)90014-0.

6. Gulbis, J.M., Kelman, Z., Hurwitz, J., O’Donnell, M., and Kuriyan, J. (1996). Structure of the C-terminal region of p21(WAF1/CIP1) complexed with human PCNA. Cell 87, 297–306. 10.1016/s0092-8674(00)81347-1.

7. Jeruzalmi, D., O’Donnell, M., and Kuriyan, J. (2001). Crystal structure of the processivity clamp loader gamma (gamma) complex of E. coli DNA polymerase III. Cell 106, 429–441. 10.1016/s0092-8674(01)00463-9.

8. Johnson, A., and O’Donnell, M. (2005). Cellular DNA replicases: components and dynamics at the replication fork. Annu Rev Biochem 74, 283–315. 10.1146/annurev.biochem.73.011303.073859.

9. Ellison, V., and Stillman, B. (1998). Reconstitution of recombinant human replication factor C (RFC) and identification of an RFC subcomplex possessing DNA-dependent ATPase activity. The Journal of biological chemistry 273, 5979–5987. 10.1074/jbc.273.10.5979.

10. Bowman, G.D., O’Donnell, M., and Kuriyan, J. (2004). Structural analysis of a eukaryotic sliding DNA clamp-clamp loader complex. Nature 429, 724–730. 10.1038/nature02585.

11. Kelch, B., Makino, D., Simonetta, K., O’Donnell, M., and Kuriyan, J. (2012). Molecular Mechanisms of DNA Polymerase Clamp Loaders. Macromolecular Crystallography, 103–114.

12. Gaubitz, C., Liu, X., Pajak, J., Stone, N.P., Hayes, J.A., Demo, G., and Kelch, B.A. (2022). Cryo-EM structures reveal high-resolution mechanism of a DNA polymerase sliding clamp loader. Elife 11. 10.7554/eLife.74175.

13. Zheng, F., Georgescu, R., Yao, N.Y., Li, H., and O’Donnell, M.E. (2022). Cryo-EM structures reveal that RFC recognizes both the 3’- and 5’-DNA ends to load PCNA onto gaps for DNA repair. Elife 11. 10.7554/eLife.77469.

14. Liu, X., Gaubitz, C., Pajak, J., and Kelch, B.A. (2022). A second DNA binding site on RFC facilitates clamp loading at gapped or nicked DNA. Elife 11. 10.7554/eLife.77483.

15. Schrecker, M., Castaneda, J.C., Devbhandari, S., Kumar, C., Remus, D., and Hite, R.K. (2022). Multistep loading of a DNA sliding clamp onto DNA by replication factor C. Elife 11. 10.7554/eLife.78253.

16. Xu, Z.Q., Jergic, S., Lo, A.T.Y., Pradhan, A.C., Brown, S.H.J., Bouwer, J.C., Ghodke, H., Lewis, P.J., Tolun, G., Oakley, A.J., and Dixon, N.E. (2024). Structural characterisation of the complete cycle of sliding clamp loading in Escherichia coli. Nat Commun 15, 8372. 10.1038/s41467-024-52623-9.

17. Landeck, J.T., Pajak, J., Norman, E.K., Sedivy, E.L., and Kelch, B.A. (2024). Differences between bacteria and eukaryotes in clamp loader mechanism, a conserved process underlying DNA replication. The Journal of biological chemistry 300, 107166. 10.1016/j.jbc.2024.107166.

18. Turner, J., Hingorani, M.M., Kelman, Z., and O’Donnell, M. (1999). The internal workings of a DNA polymerase clamp-loading machine. EMBO J 18, 771–783. 10.1093/emboj/18.3.771.

19. Johnson, A., and O’Donnell, M. (2003). Ordered ATP hydrolysis in the gamma complex clamp loader AAA+ machine. The Journal of biological chemistry 278, 14406–14413. 10.1074/jbc.M212708200.

20. Johnson, A., Yao, N.Y., Bowman, G.D., Kuriyan, J., and O’Donnell, M. (2006). The replication factor C clamp loader requires arginine finger sensors to drive DNA binding and proliferating cell nuclear antigen loading. The Journal of biological chemistry 281, 35531–35543. 10.1074/jbc.M606090200.

21. Li, H., O’Donnell, M., and Kelch, B. (2022). Unexpected new insights into DNA clamp loaders: Eukaryotic clamp loaders contain a second DNA site for recessed 5’ ends that facilitates repair and signals DNA damage: Eukaryotic clamp loaders contain a second DNA site for recessed 5’ ends that facilitates repair and signals DNA damage. Bioessays 44, e2200154. 10.1002/bies.202200154.

22. Liu, J., Zhou, Y., and Hingorani, M.M. (2017). Linchpin DNA-binding residues serve as go/no-go controls in the replication factor C-catalyzed clamp-loading mechanism. The Journal of biological chemistry 292, 15892–15906. 10.1074/jbc.M117.798702.

23. Marzahn, M.R., Hayner, J.N., Meyer, J.A., and Bloom, L.B. (2015). Kinetic analysis of PCNA clamp binding and release in the clamp loading reaction catalyzed by Saccharomyces cerevisiae replication factor C. Biochim Biophys Acta 1854, 31–38. 10.1016/j.bbapap.2014.09.019.

24. Sakato, M., Zhou, Y., and Hingorani, M.M. (2012). ATP binding and hydrolysis-driven rate-determining events in the RFC-catalyzed PCNA clamp loading reaction. J Mol Biol 416, 176–191. 10.1016/j.jmb.2011.12.018.

25. Zheng, F., Georgescu, R.E., Yao, N.Y., O’Donnell, M.E., and Li, H. (2022). DNA is loaded through the 9-1-1 DNA checkpoint clamp in the opposite direction of the PCNA clamp. Nat Struct Mol Biol 29, 376–385. 10.1038/s41594-022-00742-6.

26. Castaneda, J.C., Schrecker, M., Remus, D., and Hite, R.K. (2022). Mechanisms of loading and release of the 9-1-1 checkpoint clamp. Nat Struct Mol Biol 29, 369–375. 10.1038/s41594-022-00741-7.

27. Day, M., Oliver, A.W., and Pearl, L.H. (2022). Structure of the human RAD17-RFC clamp loader and 9-1-1 checkpoint clamp bound to a dsDNA-ssDNA junction. Nucleic Acids Res 50, 8279–8289. 10.1093/nar/gkac588.

28. Simonetta, K.R., Kazmirski, S.L., Goedken, E.R., Cantor, A.J., Kelch, B.A., McNally, R., Seyedin, S.N., Makino, D.L., O’Donnell, M., and Kuriyan, J. (2009). The mechanism of ATP-dependent primer-template recognition by a clamp loader complex. Cell 137, 659–671. 10.1016/j.cell.2009.03.044.

29. Onrust, R., Finkelstein, J., Naktinis, V., Turner, J., Fang, L., and O’Donnell, M. (1995). Assembly of a chromosomal replication machine: two DNA polymerases, a clamp loader, and sliding clamps in one holoenzyme particle. I. Organization of the clamp loader. The Journal of biological chemistry 270, 13348–13357. 10.1074/jbc.270.22.13348.

30. Yao, N., Turner, J., Kelman, Z., Stukenberg, P.T., Dean, F., Shechter, D., Pan, Z.Q., Hurwitz, J., and O’Donnell, M. (1996). Clamp loading, unloading and intrinsic stability of the PCNA, beta and gp45 sliding clamps of human, E. coli and T4 replicases. Genes Cells 1, 101–113. 10.1046/j.1365-2443.1996.07007.x.

31. Zheng, F., Georgescu, R.E., Yao, N.Y., O’Donnell, M.E., and Li, H. (2023). Structures of 9-1-1 DNA checkpoint clamp loading at gaps from start to finish and ramification on biology. Cell Rep 42, 112694. 10.1016/j.celrep.2023.112694.

32. Yao, N., Leu, F.P., Anjelkovic, J., Turner, J., and O’Donnell, M. (2000). DNA structure requirements for the Escherichia coli gamma complex clamp loader and DNA polymerase III holoenzyme. The Journal of biological chemistry 275, 11440–11450. 10.1074/jbc.275.15.11440.

33. Ason, B., Handayani, R., Williams, C.R., Bertram, J.G., Hingorani, M.M., O’Donnell, M., Goodman, M.F., and Bloom, L.B. (2003). Mechanism of loading the Escherichia coli DNA polymerase III beta sliding clamp on DNA. Bona fide primer/templates preferentially trigger the gamma complex to hydrolyze ATP and load the clamp. The Journal of biological chemistry 278, 10033–10040. 10.1074/jbc.M211741200.

34. Snyder, A.K., Williams, C.R., Johnson, A., O’Donnell, M., and Bloom, L.B. (2004). Mechanism of loading the Escherichia coli DNA polymerase III sliding clamp: II. Uncoupling the beta and DNA binding activities of the gamma complex. The Journal of biological chemistry 279, 4386–4393. 10.1074/jbc.M310430200.

35. Gaubitz, C., Liu, X., Magrino, J., Stone, N.P., Landeck, J., Hedglin, M., and Kelch, B.A. (2020). Structure of the human clamp loader reveals an autoinhibited conformation of a substrate-bound AAA+ switch. Proc Natl Acad Sci U S A 117, 23571–23580. 10.1073/pnas.2007437117.

36. Marceau, A.H., Bahng, S., Massoni, S.C., George, N.P., Sandler, S.J., Marians, K.J., and Keck, J.L. (2011). Structure of the SSB-DNA polymerase III interface and its role in DNA replication. EMBO J 30, 4236–4247. 10.1038/emboj.2011.305.

37. Tondnevis, F., Gillilan, R.E., Bloom, L.B., and McKenna, R. (2015). Solution study of the Escherichia coli DNA polymerase III clamp loader reveals the location of the dynamic psichi heterodimer. Struct Dyn 2, 054701. 10.1063/1.4927407.

38. Gomes, X.V., Gary, S.L., and Burgers, P.M. (2000). Overproduction in Escherichia coli and characterization of yeast replication factor C lacking the ligase homology domain. The Journal of biological chemistry 275, 14541–14549. 10.1074/jbc.275.19.14541.

39. Rajagopalan, M., Lu, C., Woodgate, R., O’Donnell, M., Goodman, M.F., and Echols, H. (1992). Activity of the purified mutagenesis proteins UmuC, UmuD’, and RecA in replicative bypass of an abasic DNA lesion by DNA polymerase III. Proc Natl Acad Sci U S A 89, 10777–10781. 10.1073/pnas.89.22.10777.

40. Indiani, C., Langston, L.D., Yurieva, O., Goodman, M.F., and O’Donnell, M. (2009). Translesion DNA polymerases remodel the replisome and alter the speed of the replicative helicase. Proc Natl Acad Sci U S A 106, 6031–6038. 10.1073/pnas.0901403106.

41. Indiani, C., McInerney, P., Georgescu, R., Goodman, M.F., and O’Donnell, M. (2005). A sliding-clamp toolbelt binds high- and low-fidelity DNA polymerases simultaneously. Mol Cell 19, 805–815. 10.1016/j.molcel.2005.08.011.

42. Indiani, C., Patel, M., Goodman, M.F., and O’Donnell, M.E. (2013). RecA acts as a switch to regulate polymerase occupancy in a moving replication fork. Proc Natl Acad Sci U S A 110, 5410–5415. 10.1073/pnas.1303301110.

43. Bonner, C.A., Stukenberg, P.T., Rajagopalan, M., Eritja, R., O’Donnell, M., McEntee, K., Echols, H., and Goodman, M.F. (1992). Processive DNA synthesis by DNA polymerase II mediated by DNA polymerase III accessory proteins. The Journal of biological chemistry 267, 11431–11438.

44. Wagner, J., Fujii, S., Gruz, P., Nohmi, T., and Fuchs, R.P. (2000). The beta clamp targets DNA polymerase IV to DNA and strongly increases its processivity. EMBO Rep 1, 484–488. 10.1093/embo-reports/kvd109.

45. Bloom, L.B. (2009). Loading clamps for DNA replication and repair. DNA Repair (Amst) 8, 570–578. 10.1016/j.dnarep.2008.12.014.

46. Lopez de Saro, F.J., and O’Donnell, M. (2001). Interaction of the beta sliding clamp with MutS, ligase, and DNA polymerase I. Proc Natl Acad Sci U S A 98, 8376–8380. 10.1073/pnas.121009498.

47. Becherel, O.J., Fuchs, R.P., and Wagner, J. (2002). Pivotal role of the beta-clamp in translesion DNA synthesis and mutagenesis in E. coli cells. DNA Repair (Amst) 1, 703–708. 10.1016/s1568-7864(02)00106-4.

48. Yang, W. (2003). Damage repair DNA polymerases Y. Curr Opin Struct Biol 13, 23–30. 10.1016/s0959-440x(02)00003-9.

49. Maul, R.W., Ponticelli, S.K., Duzen, J.M., and Sutton, M.D. (2007). Differential binding of Escherichia coli DNA polymerases to the beta-sliding clamp. Mol Microbiol 65, 811–827. 10.1111/j.1365-2958.2007.05828.x.

50. Kath, J.E., Chang, S., Scotland, M.K., Wilbertz, J.H., Jergic, S., Dixon, N.E., Sutton, M.D., and Loparo, J.J. (2016). Exchange between Escherichia coli polymerases II and III on a processivity clamp. Nucleic Acids Res 44, 1681–1690. 10.1093/nar/gkv1375.

51. Furukohri, A., Goodman, M.F., and Maki, H. (2008). A dynamic polymerase exchange with Escherichia coli DNA polymerase IV replacing DNA polymerase III on the sliding clamp. The Journal of biological chemistry 283, 11260–11269. 10.1074/jbc.M709689200.

52. Kath, J.E., Jergic, S., Heltzel, J.M., Jacob, D.T., Dixon, N.E., Sutton, M.D., Walker, G.C., and Loparo, J.J. (2014). Polymerase exchange on single DNA molecules reveals processivity clamp control of translesion synthesis. Proc Natl Acad Sci U S A 111, 7647–7652. 10.1073/pnas.1321076111.

53. Fijalkowska, I.J., Schaaper, R.M., and Jonczyk, P. (2012). DNA replication fidelity in Escherichia coli: a multi-DNA polymerase affair. FEMS Microbiol Rev 36, 1105–1121. 10.1111/j.1574-6976.2012.00338.x.

54. Li, N., Lam, W.H., Zhai, Y., Cheng, J., Cheng, E., Zhao, Y., Gao, N., and Tye, B.K. (2018). Structure of the origin recognition complex bound to DNA replication origin. Nature 559, 217–222. 10.1038/s41586-018-0293-x.

55. Correction for Yuan et al., Structural mechanism of helicase loading onto replication origin DNA by ORC-Cdc6. (2021). Proc Natl Acad Sci U S A 118. 10.1073/pnas.2025413118.

56. Neylon, C., Kralicek, A.V., Hill, T.M., and Dixon, N.E. (2005). Replication termination in Escherichia coli: structure and antihelicase activity of the Tus-Ter complex. Microbiol Mol Biol Rev 69, 501–526. 10.1128/MMBR.69.3.501-526.2005.

57. Simonsen, S., Sogaard, C.K., Olsen, J.G., Otterlei, M., and Kragelund, B.B. (2024). The bacterial DNA sliding clamp, beta-clamp: structure, interactions, dynamics and drug discovery. Cell Mol Life Sci 81, 245. 10.1007/s00018-024-05252-w.

58. Moolman, M.C., Krishnan, S.T., Kerssemakers, J.W., van den Berg, A., Tulinski, P., Depken, M., Reyes-Lamothe, R., Sherratt, D.J., and Dekker, N.H. (2014). Slow unloading leads to DNA-bound beta2-sliding clamp accumulation in live Escherichia coli cells. Nat Commun 5, 5820. 10.1038/ncomms6820.

59. Lazowski, K., Woodgate, R., and Fijalkowska, I.J. (2024). Escherichia coli DNA replication: the old model organism still holds many surprises. FEMS Microbiol Rev 48. 10.1093/femsre/fuae018.

60. Fernandez-Leiro, R., Conrad, J., Scheres, S.H.W., and Lamers, M.H. (2015). cryo-EM structures of the E. coli replicative DNA polymerase reveal its dynamic interactions with the DNA sliding clamp, exonuclease and τ. eLife 4, e11134. 10.7554/eLife.11134.

61. Reyes-Lamothe, R., Sherratt, D.J., and Leake, M.C. (2010). Stoichiometry and architecture of active DNA replication machinery in Escherichia coli. Science 328, 498–501. 10.1126/science.1185757.

62. McInerney, P., Johnson, A., Katz, F., and O’Donnell, M. (2007). Characterization of a triple DNA polymerase replisome. Mol Cell 27, 527–538. 10.1016/j.molcel.2007.06.019.

63. Kelman, Z., Yuzhakov, A., Andjelkovic, J., and O’Donnell, M. (1998). Devoted to the lagging strand-the subunit of DNA polymerase III holoenzyme contacts SSB to promote processive elongation and sliding clamp assembly. EMBO J 17, 2436–2449. 10.1093/emboj/17.8.2436.

64. Kelman, Z., Yao, N., and O’Donnell, M. (1995). Escherichia coli expression vectors containing a protein kinase recognition motif, His6-tag and hemagglutinin epitope. Gene 166, 177–178. 10.1016/0378-1119(95)00556-7.

65. Studwell, P.S., and O’Donnell, M. (1990). Processive replication is contingent on the exonuclease subunit of DNA polymerase III holoenzyme. The Journal of biological chemistry 265, 1171–1178.

66. Kelman, Z., Naktinis, V., and O’Donnell, M. (1995). Radiolabeling of proteins for biochemical studies. Methods Enzymol 262, 430–442. 10.1016/0076-6879(95)62034-6.

67. Mastronarde, D.N. (2003). SerialEM: A Program for Automated Tilt Series Acquisition on Tecnai Microscopes Using Prediction of Specimen Position. Microscopy and Microanalysis 9, 1182–1183.

68. Punjani, A., Rubinstein, J.L., Fleet, D.J., and Brubaker, M.A. (2017). cryoSPARC: algorithms for rapid unsupervised cryo-EM structure determination. Nat Methods 14, 290–296. 10.1038/nmeth.4169.

69. Scheres, S.H. (2012). RELION: implementation of a Bayesian approach to cryo-EM structure determination. J Struct Biol 180, 519–530. 10.1016/j.jsb.2012.09.006.

70. Zheng, S.Q., Palovcak, E., Armache, J.P., Verba, K.A., Cheng, Y., and Agard, D.A. (2017). MotionCor2: anisotropic correction of beam-induced motion for improved cryo-electron microscopy. Nat Methods 14, 331–332. 10.1038/nmeth.4193.

71. Rohou, A., and Grigorieff, N. (2015). CTFFIND4: Fast and accurate defocus estimation from electron micrographs. J Struct Biol 192, 216–221. 10.1016/j.jsb.2015.08.008.

72. Bepler, T., Morin, A., Rapp, M., Brasch, J., Shapiro, L., Noble, A.J., and Berger, B. (2019). Positive-unlabeled convolutional neural networks for particle picking in cryo-electron micrographs. Nat Methods 16, 1153–1160. 10.1038/s41592-019-0575-8.

73. Asarnow, D., Palovcak, E., and Cheng, Y. (2019). UCSF pyem v0. 5. Zenodo https://doi.org/10.5281/zenodo *3576630*.

74. Pettersen, E.F., Goddard, T.D., Huang, C.C., Couch, G.S., Greenblatt, D.M., Meng, E.C., and Ferrin, T.E. (2004). UCSF Chimera--a visualization system for exploratory research and analysis. J Comput Chem 25, 1605–1612. 10.1002/jcc.20084.

75. Goddard, T.D., Huang, C.C., Meng, E.C., Pettersen, E.F., Couch, G.S., Morris, J.H., and Ferrin, T.E. (2018). UCSF ChimeraX: Meeting modern challenges in visualization and analysis. Protein Sci 27, 14–25. 10.1002/pro.3235.

76. Emsley, P., Lohkamp, B., Scott, W.G., and Cowtan, K. (2010). Features and development of Coot. Acta Crystallogr D Biol Crystallogr 66, 486–501. 10.1107/S0907444910007493.

77. Adams, P.D., Afonine, P.V., Bunkoczi, G., Chen, V.B., Davis, I.W., Echols, N., Headd, J.J., Hung, L.W., Kapral, G.J., Grosse-Kunstleve, R.W., et al. (2010). PHENIX: a comprehensive Python-based system for macromolecular structure solution. Acta Crystallogr D Biol Crystallogr 66, 213–221. 10.1107/S0907444909052925.

78. Zheng, F., Georgescu, R.E., Li, H., and O’Donnell, M.E. (2020). Structure of eukaryotic DNA polymerase delta bound to the PCNA clamp while encircling DNA. Proc Natl Acad Sci U S A 117, 30344–30353. 10.1073/pnas.2017637117.

79. Sanchez-Garcia, R., Gomez-Blanco, J., Cuervo, A., Carazo, J.M., Sorzano, C.O.S., and Vargas, J. (2021). DeepEMhancer: a deep learning solution for cryo-EM volume post-processing. Commun Biol 4, 874. 10.1038/s42003-021-02399-1.

80. He, J., Li, T., and Huang, S.Y. (2023). Improvement of cryo-EM maps by simultaneous local and non-local deep learning. Nat Commun 14, 3217. 10.1038/s41467-023-39031-1.

81. Schwab, J., Kimanius, D., Burt, A., Dendooven, T., and Scheres, S.H.W. (2024). DynaMight: estimating molecular motions with improved reconstruction from cryo-EM images. Nat Methods 21, 1855–1862. 10.1038/s41592-024-02377-5.

82. Asarnow, D., Palovcak, E., and Cheng, Y. (2019). UCSF pyem v0. 5. Zenodo 3576630. 10.5281/zenodo.3576630.

83. Georgescu, R.E., Kim, S.-S., Yurieva, O., Kuriyan, J., Kong, X.-P., and O’Donnell, M. (2008). Structure of a Sliding Clamp on DNA. Cell 132, 43–54. 10.1016/j.cell.2007.11.045.

84. Jamali, K., Kall, L., Zhang, R., Brown, A., Kimanius, D., and Scheres, S.H.W. (2024). Automated model building and protein identification in cryo-EM maps. Nature 628, 450–457. 10.1038/s41586-024-07215-4.

85. Chen, V.B., Arendall, W.B., 3rd, Headd, J.J., Keedy, D.A., Immormino, R.M., Kapral, G.J., Murray, L.W., Richardson, J.S., and Richardson, D.C. (2010). MolProbity: all-atom structure validation for macromolecular crystallography. Acta Crystallogr D Biol Crystallogr 66, 12–21. 10.1107/S0907444909042073.

86. Pettersen, E.F., Goddard, T.D., Huang, C.C., Meng, E.C., Couch, G.S., Croll, T.I., Morris, J.H., and Ferrin, T.E. (2021). UCSF ChimeraX: Structure visualization for researchers, educators, and developers. Protein Sci 30, 70–82. 10.1002/pro.3943.

87. Oakley, A.J., Prosselkov, P., Wijffels, G., Beck, J.L., Wilce, M.C., and Dixon, N.E. (2003). Flexibility revealed by the 1.85 A crystal structure of the beta sliding-clamp subunit of Escherichia coli DNA polymerase III. Acta Crystallogr D Biol Crystallogr 59, 1192–1199. 10.1107/s0907444903009958.

88. Kimanius, D., Jamali, K., Wilkinson, M.E., Lövestam, S., Velazhahan, V., Nakane, T., and Scheres, S.H.W. (2023). Data-driven regularisation lowers the size barrier of cryo-EM structure determination. bioRxiv, 2023.2010.2023.563586. 10.1101/2023.10.23.563586.

89. Stewart, J., Hingorani, M.M., Kelman, Z., and O’Donnell, M. (2001). Mechanism of beta clamp opening by the delta subunit of Escherichia coli DNA polymerase III holoenzyme. The Journal of biological chemistry 276, 19182–19189. 10.1074/jbc.M100592200.

90. Edgar, R.C. (2004). MUSCLE: multiple sequence alignment with high accuracy and high throughput. Nucleic Acids Res 32, 1792–1797. 10.1093/nar/gkh340.

